# Effects of single cage housing on stress, cognitive and seizure parameters in the rat and mouse pilocarpine models of epilepsy

**DOI:** 10.1101/214528

**Authors:** H Manouze, A Ghestem, V Poillerat, M Bennis, S Ba-M’hamed, JJ Benoliel, C Becker, C Bernard

## Abstract

Many experimental approaches require housing rodents in individual cages, including in epilepsy research. However, rats and mice are social animals; and individual housing constitutes a stressful situation. The goal of the present study was to determine the effects of individual housing as compared to conditions maintaining social contact on stress markers and epilepsy. Control male mice socially housed during pretest and then transferred to individual cages for six weeks displayed anhedonia, increased anxiety and biological markers of stress as compared to pretest values or mice kept socially housed during six weeks. Pilocarpine-treated mice housed together showed increased levels of anhedonia, anxiety and stress markers as well as decreased cognitive performance as compared to the control group. The differences were more significant in pilocarpine-treated mice housed individually. Anxiety correlated linearly with cognitive performance and stress markers independently of the experimental conditions. In the male rat pilocarpine model, seizures were sixteen times more frequent in singly housed animals as compared to animals kept in pairs. Daily interactions with an experimenter in otherwise singly housed animals was sufficient to produce results identical to those found in animals kept in pairs. We propose that social isolation produces a severe phenotype in terms of stress and seizure frequency as compared to animals maintaining social contact (at least in these two models), a factor that needs to be taken into account for data interpretation, in particular for preclinical studies.

**Significance Statement:** Many experimental approaches require housing rodents in individual cages, a stressful condition for social animals, even in an enriched environment context. Using the pilocarpine model of epilepsy in rats and mice, we report that singly housing animals develop a more severe phenotype in terms of stress and epilepsy as compared to animals maintaining social contact. We propose that social isolation adds a degree of complexity for the interpretation of data, which may be particularly relevant for preclinical studies.

## Introduction

Social isolation (single housing) and barren environment are stressful to rodents; for a recent review cf. (Arakawa, 2018; Mumtaz et al., 2018). Social isolation of rodents after weaning is often used as an experimental manipulation to model early life stress in humans. It produces anxiety/depressive-like behavior and cognitive deficits and favors the emergence of pathological traits such as addictive behavior (Bianchi et al., 2006; Brenes et al., 2008; Ros-Simo and Valverde, 2012). These behavioral alterations are associated with biological modifications including oxidative stress, the production of stress hormones and inflammatory cytokines (Moller et al., 2013; Krugel et al., 2014; Shao et al., 2015; Butler et al., 2016). Social isolation performed in adult animals also results in behavioral and endocrinological alterations (Berry et al., 2012; Krugel et al., 2014; Ieraci et al., 2016; Filipovic et al., 2017). However, a general understanding of the consequences of social isolation is difficult to extract from these studies because they use different types of isolation, period and age of isolation, behavioral and molecular protocols, leading to conflicting reports (Arakawa, 2018). Since all studies report that social isolation does produce biological alterations, many regulation agencies recommend social housing and environmental enrichment to improve the welfare of rats and mice in animal facilities (Lidster et al., 2016). However, single housing of rodents may be imposed by the experimental protocol to obtain accurate measurements of biological parameters in individual animals, such as in the fields of toxicology, drug addiction, and neurological disorders. Experimental epilepsy is a typical example. In order to precisely assess seizures with continuous video-EEG recordings, instrumented animals are singly housed to prevent the severing of EEG wires and aggressive behavior from others and to clearly identify individuals on the video. Several studies have shown that environmental enrichment has positive effects on seizure frequency or severity in experimental models (Morelli et al., 2014; Kotloski and Sutula, 2015; Dezsi et al., 2016; Vrinda et al., 2017), but the effect of social isolation starting from the beginning of the experimental procedure in the presence of environmental enrichment has not been assessed.

There exist many types of experimental models of epilepsy in different rodent species and strains (Levesque et al., 2016; Loscher, 2017; Becker, 2018). The effects of social isolation should be studied for each model, as results may be model-, strain- and species-specific. For example, Wistar and Sprague Dawley rats display different phenotypes in terms of depression-like profile and cognitive deficits in the kainic acid and pilocarpine models (Inostroza et al., 2011; Inostroza et al., 2012). Here we focus on the widely used pilocarpine model of experimental epilepsy in adult male Swiss mice and Wistar rats (Levesque et al., 2016). Numerous studies show a dysregulation of the HPA axis in patients and in experimental epilepsy, which is model-dependent (Maguire and Salpekar, 2013; Wulsin et al., 2016; Mumtaz et al., 2018). Since stress also results in the activation of the HPA axis, we reasoned that social isolation in experimental models of epilepsy might produce a strong HPA axis response via the contributions of both isolation- and epilepsy-induced stress (Gunn and Baram, 2017). Social isolation during the neonatal or juvenile periods increases seizure susceptibility and produces a behavioral phenotype (Lai et al., 2006; Amiri et al., 2017), but the effects of social isolation in adult experimental models of epilepsy have not been investigated. The issue is important as most mechanistic, pharmacological and interventional studies use recordings of singly housed animals. We hypothesized that the combination of both isolation x epilepsy factors might exacerbate the epilepsy phenotype as compared to animals maintaining social interactions. Clinical studies support this hypothesis. Stressful life events are associated with an increased risk of seizure occurrence in patients with epilepsy (Baldin et al., 2017; Kotwas et al., 2017). In some patients, the fear to have a seizure and social isolation due to stigmatization may increase the allostatic load, with a direct impact on seizure frequency (Kotwas et al., 2017). We thus assessed the consequences of animal housing on stress response and epilepsy severity, acting on a single experimental variable: social interaction (environmental enrichment was provided). We also propose an alternate solution to maintain social contact in singly housed animals, to decrease their stress level. We hope to foster discussions on how to interpret results (including negative preclinical studies) obtained in different housing conditions.

## METHODS

### Animals

We used male Wistar rats (200 to 250g; Charles Rivers Laboratories, Les Oncins, France), aged nine weeks (at their arrival in the laboratory). Swiss male mice (8 weeks old) were obtained from the animal husbandry of the Faculty of Sciences, Cadi Ayyad University, Marrakech (Morocco). The animals were kept under controlled environmental conditions (23±1 °C; night-day cycle (12 h-12 h)) with *ad libitum* access to food and water. Zeitgeber (ZT) 0 was at 7:30 am (the time when the light was switched on in the animal facility). All procedures were conducted per approved institutional protocols, and with the provisions for animal care and use prescribed in the scientific procedures on living animals, European Council Directive: EU2010/63 and by the Council Committee of Research Laboratories of the Faculty of Sciences, Cadi Ayyad University, Marrakech. All efforts were made to minimize any animal suffering.

### Experimental design

#### In mice

The experimental protocol is shown in Figure 1A. Swiss male mice were housed in groups of 3 per cage. In order to validate stress in the same mice throughout the experimental protocol, one mouse from each litter was randomly picked, ear-marked and returned to the same cage with littermates. They were left together (group-housed) during one week without experimental intervention to allow adaptation to the housing environment. After that, the ear-marked mouse of each litter was subjected to different behavioral tests in order to obtain their pretest levels in the social condition. Sucrose preference was assessed every day for one week. At the end of the week, we evaluated anxiety levels (Day-3: elevated plus maze test – EPM) and memory functions (Day -2-1: novel object recognition test -NOR). In this experimental protocol, we consider two independent variables: housing (social and isolation) and treatment (control and epilepsy), thus defining four groups of animals. After establishing pretest levels, cages were randomly assigned to two groups to receive pilocarpine injection to trigger status epilepticus (Pilo group– experimental procedure described hereafter) or saline (Non-pilo group). Both groups were further divided into two groups: social condition (SC, n =18, 3/cage) and isolated condition (IC, n = 6, 1/cage). For the isolated condition, ear-marked mice were taken away from their home-cages and singly housed in cages identical to their home-cages. For the social condition, the ear-marked mice remained in their home-cages with littermates. We refer to the time following group assignment as the posttest period. Four weeks following the beginning of the posttest period, mice already evaluated during the pretest period were again evaluated for anhedonia (Day 28-35), EPM (Day 36) and NOR (37-38). Behavioral tasks (EPM and NOR) were recorded and analyzed using Ethovision®XTNoldus 8.5 video tracking program (Noldus, Netherlands) connected to a video camera (JVC). At the end of each behavioral session, apparatuses were cleaned with a 75% ethanol solution to remove any of odor or trace. The behavioral tests were performed between 8:00 and 12:00 a.m. during the light cycle to avoid the circadian-related fluctuation in the performance of the mice. Before the beginning of behavioral tests, the animals were transferred to the testing room in their home cages and left there to habituate for 60 min.

**Figure 1:**
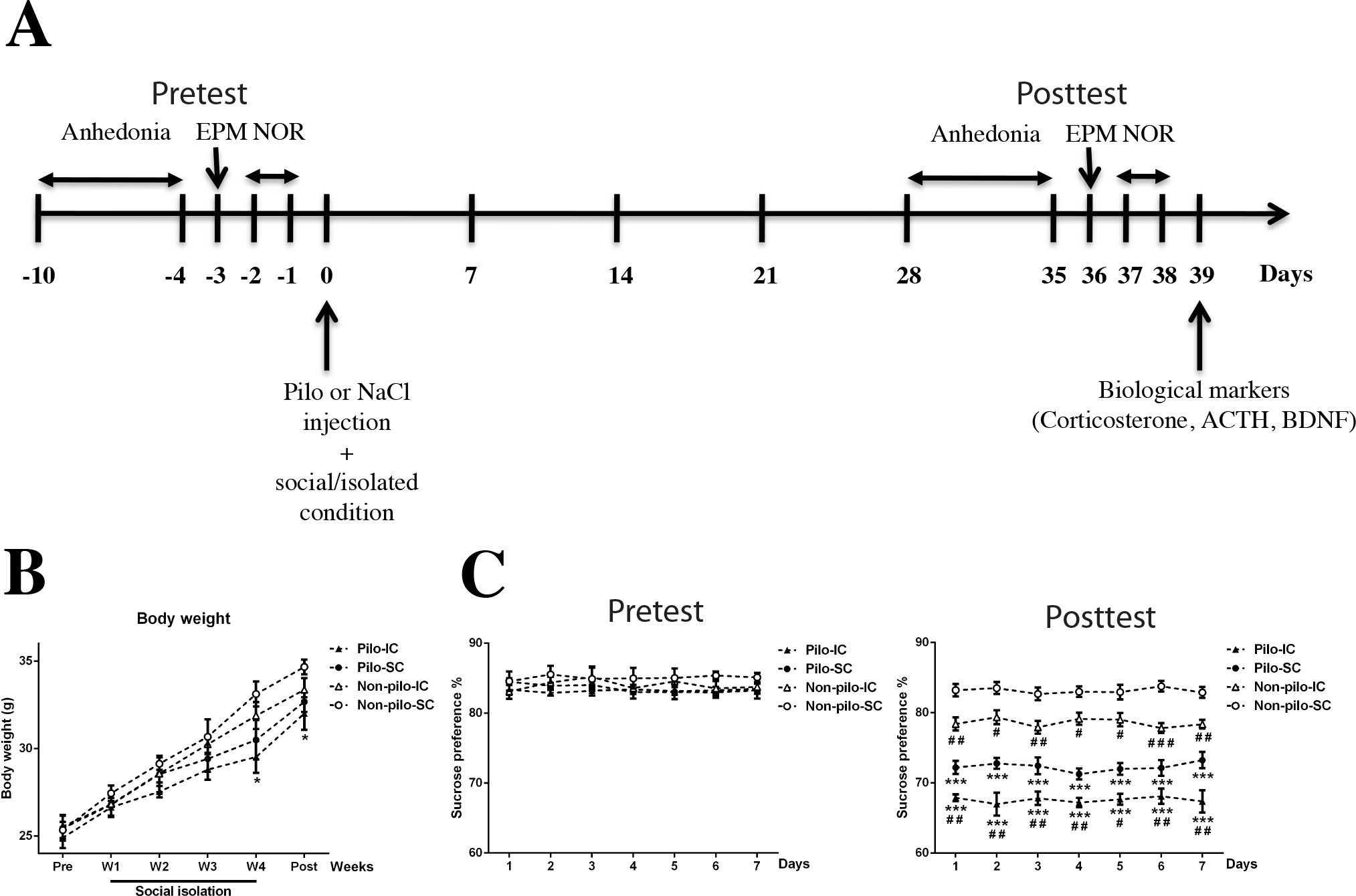
Effect of social isolation on body weight and anhedonia in control and epileptic mice. **(A)** Experimental protocol. In pretest condition, up to day 0, mice were socially housed and evaluated for anhedonia, anxiety (EPM) and novel object recognition (NOR) test. Tested mice were then assigned randomly to different groups: Non-pilo social condition (SC), i.e. control like before D0, Non-pilo isolated condition (IC), i.e. the mice were transferred to an individual cage, Pilo SC, the mice received pilocarpine to induce status epilepticus, which triggers epileptogenesis, the process leading to the occurrence of spontaneous seizures, and Pilo IC. Four weeks following status epilepticus (posttest conditions), the same mice were evaluated for anhedonia, EPM and NOR tests. At the end of the experimental protocol, animals were killed to measure corticosterone, ACTH, and BDNF. **(B)** Evolution of averaged body weight over six weeks for the four groups. The Pilo IC groups gained less weight than the other groups. Data are mean ± SEM. \**p*<0.05 in comparison with the control non-Pilo SC group. (**C**) Sweet water consumption was identical in all mice in pretest conditions (splitting them on their future group assignment). As compared to the control group (Non-pilo SC), the Non-pilo IC showed anhedonia, which was further increased in the Pilo-SC and exacerbated in the Pilo-IC group. Data are mean ± SEM. \*\*\**p*< 0.001 in comparison with the non-Pilo SC group; ^#^*p*<0.05, ^##^*p*<0.01 and ^###^*p*<0.001 in comparison with the corresponding social condition group.

At the end of the experimental protocol, we measured ACTH, CORT and BDNF levels as in rats (described below).

#### In rats

The experimental protocol is shown in Figure 7A. Animals were received in sets of 4 from the same litter from the vendor. Two main groups of animals were used: animals with spontaneous seizures following pilocarpine-induced status epilepticus (pilo group – experimental procedure described hereafter) and control animals (Non-pilo group). Pilo and Non-pilo animals were further divided into three groups:

- Isolated group: rats were singly housed and were not handled during the experimental period except for cage cleaning and body weight measurement once a week.
- Handled group: rats were singly housed and were handled daily until the end of the experiment (see below).
- Paired group: rats were kept in buddy pairs with social interaction upon arrival in the animal facility.

Handling was performed twice daily (ZT2 and ZT9) throughout the experimental period by the same experimenter. Briefly, each handling session consisted of stroking animals for 1 minute each in their cage. Then, each rat was gently handled by experimenter’s hands (without wearing gloves), while being softly stroked from the head to the tail for 2 minutes. Finally, the rats were placed back in their home cage and fed by the experimenter for 2 minutes. Isolated and paired groups were left undisturbed, except for weekly cage cleaning and body weighing.

Animals from the Non-pilo group were randomized to the three housing groups when received. Animals from the Pilo group were randomized after pilocarpine-induced status epilepticus. Since we do not know the stress history of the animals when we receive them, randomization decreases this confounding factor.

#### Sample size

A preliminary analysis made on isolated (n=4) and handled (n=4) pilo groups for seizure frequency allowed us to perform a power analysis. We determined that n=3 per group was enough. The experimental procedure in the pilo groups was conducted twice. The first and second group of simultaneously recorded animals being composed of (n=4 isolated, n=4 handled, n=6 paired) and (n=3 isolated, n=4 handled, n=6 paired), respectively. In the paired groups, only one animal was instrumented. Seizure analysis was thus performed in 6 animals. Results were similar in the two sets, and data were pooled together.

For the mice experiments, the number of subjects was defined based on the study design. Preliminary analysis was conducted with n=3 for the social condition and n=3 for the isolated condition where the results were non-significant. In this case, a second study was done with a sample size of 3 per group.

#### Status epilepticus induction and electrode implantation

In rats, status epilepticus (SE) was induced by a single i.p injection of pilocarpine (Pilo) 320 mg/kg, one week after receiving the animals from the vendor. In mice, we used repeated 100 mg/kg i.p. injections every 20 min until SE onset 17 days after receiving the animals (one week for habituation and ten days for pretest of behavioral tasks). To reduce peripheral effects, animals were pre-treated with Methyl-scopolamine (1mg/kg) 30 min prior to Pilo injection. SE was stopped by diazepam (10 mg/kg i.p., twice within a 15 min interval) after 60 min and 90 min of SE; respectively. At the end of these injections, mice and rats were hydrated with saline (2ml i.p. twice within two h) and fed with a porridge made of soaked pellets, until they resumed normal feeding behavior. All drugs were obtained from Sigma.

In rats, four weeks following SE, the telemetry implant was surgically inserted intraperitoneally under anesthesia (ketamine [1 mg/ kg]/xylasine [0.5 mg/kg] i.p) and connected to screws on the surface of the brain by two electrodes; one above the cortex (4.0 mm anteroposterior, 2.0 mm mediolaterally, compared with bregma), the second, above the cerebellum as reference. The EEG signal was transmitted by radio to a data acquisition and processing system (DSI). In the paired group, both animals developed epilepsy, but only one rat was equipped with the telemetry system as two animals cannot be recorded simultaneously with EEG transmitters in the same cage, while the other was monitored with video only. Animals were left to recover during one week before switching on the transmitter.

#### Monitoring of spontaneous recurrent seizures

Continuous (24/7) EEG recordings started in the 6th week after SE, a period sufficient to reach stability in seizure frequency (Williams et al., 2009), and were stopped at week 10. We verified that animals displayed stable seizure activity, quantifying seizure frequency during each successive week. Spontaneous recurrent seizures were detected and quantified using both visual inspections of the EEG and a semi-automatic way using Clampfit 10.2. All detected seizures were verified and reconfirmed using “NeuroScore” software. We used the following spike detection parameters: 300 µV amplitude threshold with a rejection value of 4 mV, 1 ms minimum duration, 300 ms maximum duration, 1 ms minimum spike interval, 1 s maximum spike interval, 2 s minimum spike train duration, 10 minimum spike number. All detected events were then visually inspected. These conservative parameters led to the detection of false positives, which were removed manually. In five animals, the entire EEG recording was visually inspected to confirm that no seizures escaped the semi-automatic detection (no false negative). Video recordings were performed during the light phase in the animal facility with the DSI system. They were used to assess seizure severity according to Racine’s scale after their electrophysiological detection. Rats kept in pairs never had spontaneous seizures simultaneously, which allowed a correct assessment of seizure severity of the EEG monitored animal. Finally, keeping animals in pairs did not prevent/alter continuous EEG recordings in the equipped animals (no loss of signal).

#### Behavioral and biological parameters

Body weights of mice and rats were measured weekly at ZT 2.

#### Sucrose consumption test

In mice, the baseline for sucrose consumption was assessed every day for one week, seven days after their arrival in the laboratory. The sucrose preference was calculated at the 5th week following the separation and treatment. In rats, sucrose consumption was assessed at week four and week eight following animal reception. Sucrose and water intakes were measured daily at ZT 2.

Briefly, animals were given a free choice between two bottles, one with 1% sucrose solution and another with tap water. The location of the bottles was alternated every day to prevent possible effects of side preference in drinking behavior. The consumption of water and sucrose solution was estimated by weighting the bottles. For the paired, the volume of sweet water consumed by rat was taken as the total consumed volume divided by 2. For the social groups, the volume of sweet water consumed by mouse was taken as the total consumed volume divided by the total number of animals at each time. Sweet water consumption corresponds to sucrose preference, which is calculated as a percentage of the volume of sucrose intake over the total volume of fluid intake using the following equation:

Sucrose preference= V (sucrose solution)/ (V (sucrose solution) +V (water)) X100%.

#### Elevated plus maze test

The elevated plus maze has been described as a simple method for assessing anxiety-like behavior in rodents (Pellow et al., 1985; Lapiz-Bluhm et al., 2008). The elevated plus maze consisted of four arms (two open without walls and two enclosed arms by 15 cm high walls) 50 cm long and 5 cm wide, which were joined at a square central area (5 × 5 cm) to form a plus sign. The maze floor and the side/end walls of the enclosed arms were made of clear Plexiglas. The room illumination of the elevated plus maze apparatus was under an approximate brightness of 200 lux. Briefly, the test consisted of placing the mouse gently in the central arena of the elevated plus maze, facing the junction of an open and closed arm. The mouse was allowed to freely explore the maze for 5 min while the duration and frequency of entries into open arms were recorded. Anxiety index (Rao and Sadananda, 2016) was calculated as: 1 - ((open-arm time/total time) + (open-arm entries/total entries))/2.

#### Novel object recognition test

The novel object recognition test is widely used to evaluate object recognition memory in rodents (Ennaceur and Aggleton, 1994; Reger et al., 2009; Gaskin et al., 2010). The apparatus consisted of an open field (50× 50 × 50 cm high) made of Plexiglas with the inside painted matt black. The objects to be discriminated were available as three plastic objects. 24h before testing, mice were first habituated in the open field arena in the absence of any object for 10 min. During the training trial, two identical objects (approximately 10 cm) were placed in the back corner of the box. The mouse was then positioned at the midpoint of the wall opposite to the objects and the total time spent exploring the two objects was recorded for 10 min. During the test trial (60 min after training trial), one object used during training was replaced with a novel object and both of them were placed in the middle of the back wall. The animal was then allowed to explore freely for 5 min and the time spent exploring each object was recorded. The discrimination between the novel and familiar objects was calculated by the discrimination index [(time spent exploring the novel object - time spent exploring the familial object) / total time x 100)].

#### ACTH, corticosterone and BDNF levels

Isolated, handled and paired rats in the Non-pilo group were killed by decapitation after light anesthesia with isoflurane at the beginning of the 8th week following the separation at ZT4 and the 6th weeks after the separation and treatment at ZT4 for Pilo and Non-pilo mice. Animals were decapitated in a quiet separate room, one by one, with the bench cleaned between animals. Trunk blood was collected in less than 5 sec after decapitation in a dry tube and an EDTA tube in order to obtain serum and plasma samples, respectively. The plasma was prepared by a 15 min centrifugation at 1600 g, 4°C. The ACTH concentration was determined according to the manufacturer’s instructions (Clinisciences, France). For corticosterone and BDNF levels, blood was centrifuged at 3500 g for 10 min at 4°C, and the serum was stored at 80°C until used. Corticosterone and BDNF concentrations were determined according to the manufacturer’s instructions (Coger, Promega, France).

#### Statistical analysis

The effect on body weight and anhedonia was measured using the ANOVA with repeated measures followed by Bonferroni *post-hoc* analysis using Sigma Plot 11.0 software.

For estimation based on confidence intervals, we directly introduced the raw data in https://www.estimationstats.com/ and downloaded the results and graphs. The mean difference for two comparisons is shown with Cumming estimation plot. The raw data is plotted on the upper axes. For each group, summary measurements (mean ± standard deviation) are shown as gapped lines. Each mean difference is plotted on the lower axes as a bootstrap sampling distribution. Five thousand bootstrap samples were taken; the confidence interval was bias-corrected and accelerated. Mean differences are depicted as dots; 95% confidence intervals are indicated by the ends of the vertical error bars. In order to measure the effect size, we used unbiased *Cohen’s d* (also known as standardized mean difference). We also provide *P* value(s) the likelihood(s) of observing the effect size(s), *if the null hypothesis of zero difference is true*, using the test mentioned in the text.

#### Data sharing

All raw data is stored on our NAS server and available upon request.

## RESULTS

### Effect of social isolation on weight gain and anhedonia in control and pilo mice

Repeated measures ANOVA showed a significant effect of time (F_(5,100)_=128.08, *p*<0.0001a) on body weight evolution; while the experimental group (F_(3,20)_ =2.23, *p*=0.11a) and the interaction of group and age (F_(15,100)_ =1.18, *p*=0.29a) had no significant effect (Figure 1B). *Post-hoc* analysis showed that the Pilo-IC group gained significantly less weight than the Non-pilo-SC group (*p*<0.05) at weeks four and five. There was no significant difference between all groups in body weight before social isolation until the 3^rd^ week of isolation.

We used the sucrose preference test to assess anhedonia (lack of interest in rewarding stimuli). The typical behavior of animals is a bias toward the sweetened drink. Lack of preference for the sweetened drink indicates anhedonia, which signs stress-related disorders (Gold, 2015). The ANOVA repeated measures showed an effect of group on sweet water consumption during the pretest period (F_(3,20)_=8.33, *p*<0.001^b^) and the posttest period (F_(3,20)_=295.85, *p*<0.0001^c^) with no time effect in both periods (pretest: F_(6,120)_=0.20, *p*=0.97b and posttest: F_(6,120)_=0.13, *p*=0.99^c^) (Figure 1C). During the pretest period, the *post-hoc* analysis did not show a significant difference in sucrose preference between the different groups.

During the posttest period, the *post-hoc* analysis revealed that sweet water consumption was significantly lower by 15% and 20% in the Pilo-SC and Pilo-IC groups than in the Non-pilo-SC and Non-pilo-IC groups (*p*<0.001). Both Non-pilo-IC and Pilo-IC groups had a significant decrease in sucrose preference as compared to Non-pilo-SC (*p*<0.05, *p*<0.01 and *p*<0.001) and Pilo-SC (*p*<0.05 and *p*<0.01) groups, respectively (Figure 1C). Therefore, social isolation induces anhedonia in the control (Non-pilo) group. The Pilo group socially housed also displayed anhedonia as compared to the control group. The anhedonia phenotype was further increased in the socially isolated Pilo group.

In the following, we continue to use statistical inference analysis (null hypothesis significance testing) with *P* values, but we also include estimation based on confidence intervals (CI), which provides better insight into the structure and interpretation of the results. An estimation based on confidence intervals would have been too cumbersome for the weight and anhedonia data.

### Social isolation increases anxiety levels in control and pilo mice

We then tested animals for anxiety-like behavior. All groups were tested before experimental manipulation (pretest) and again afterward (posttest). As expected, anxiety levels at pretest were very similar across groups, with each group mean within 3% of the others (Figure 2A, compare Pre scores across groups). Thus, the random assignment did not induce a bias. There was notable diversity, though, in pretest scores, with some animals scoring relatively low (0.3) and others relatively high (0.45). These differences, however, were not clearly predictive of posttest scores (*r* = 0.04 95% CI[-0.37,0.43], *p* = 0.87).

**Figure 2:**
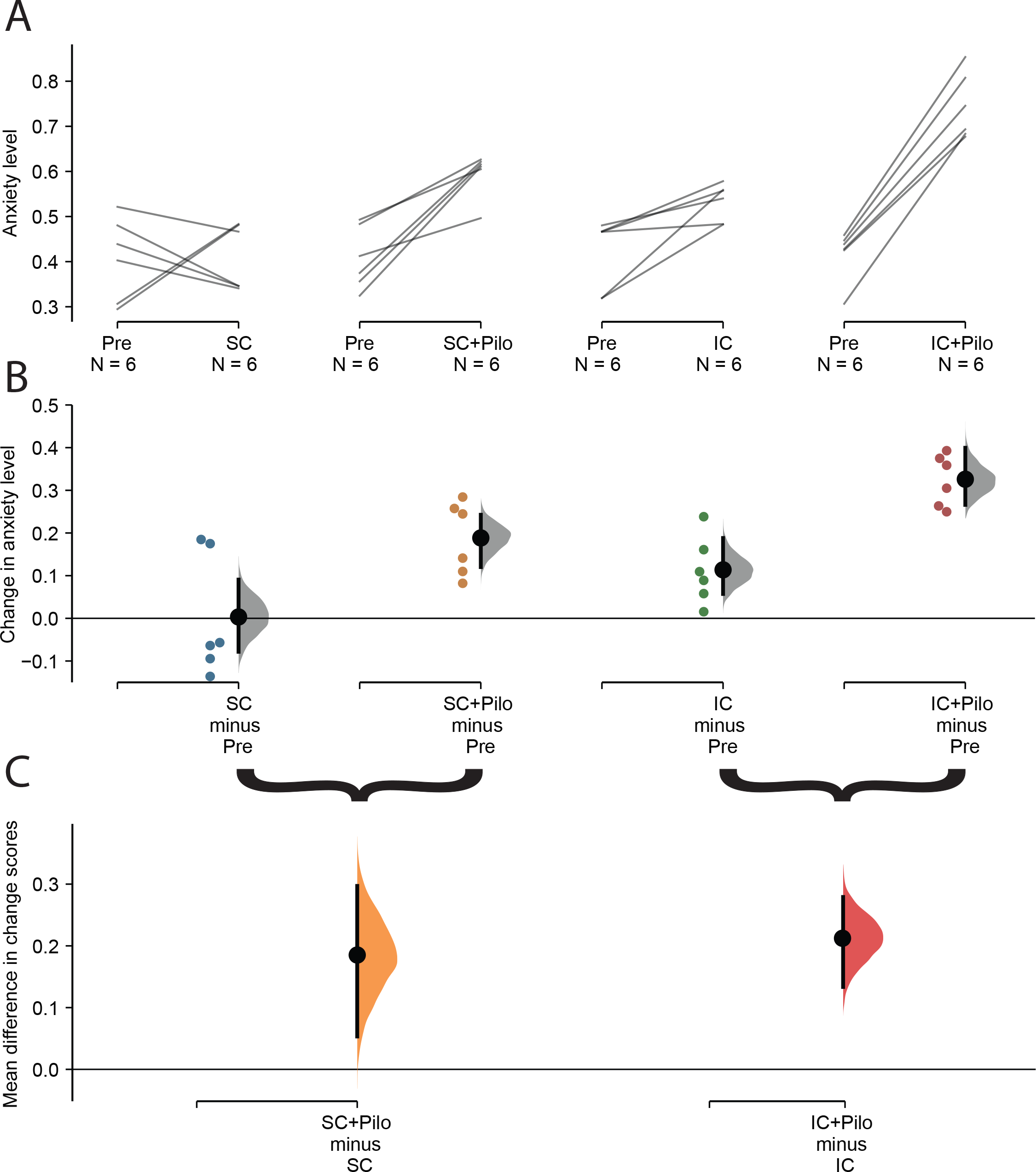
Effect of social isolation on anxiety levels in control and epileptic mice. The paired mean difference for four comparisons between posttest (Non-pilo SC, Pilo SC, Non-pilo IC, and Pilo IC) and pretest (Pre) conditions are shown in the Cumming estimation plot. The raw data is plotted on the upper axes; each paired set of observations is connected by a line (**A**). On the lower axis (**B**), each paired mean difference is plotted as a bootstrap sampling distribution. Mean differences are depicted as black dots; 95% confidence intervals are indicated by the ends of the vertical error bars. Colored dots correspond to the posttest-pretest differences for each mouse. Although anxiety levels appear similar in average in pretest conditions, we note a wide dispersion of the raw data, i.e. individual animals can have very different levels of anxiety (compare pretest values in panel A). We also note the apparent existence of a bimodal distribution. **(B)** There was no change in anxiety level in the control group (Non-pilo mice maintained in social housing) although individuals displayed variability (first panel from the left). Social isolation in Non-pilo animals (third panel from the left) significantly increased anxiety levels as compared to pretest. The changes in anxiety levels were more extensive in the Pilo-SC group (second panel) and most extensive in the Pilo-IC group (fourth panel). **(C)** Each mean difference in changes in anxiety score is plotted as a bootstrap sampling distribution. Mean differences are depicted as dots; 95% confidence intervals are indicated by the ends of the vertical error bars. Epilepsy (in Pilo mice) produced the same increase in anxiety levels in SC and IC conditions (although the IC group starts from higher anxiety levels as compared to the SC group). This means that the effect of epilepsy in isolated mice on anxiety levels is a blend of the effect of epilepsy and isolation.

For animals maintaining social housing during the posttest period, there was no average change in anxiety index scores (Mean difference score (*M*_diff_) = 0.00 95% CI[-0.08, 0.09], *d* = 0.0 95% CI[-1.2, 1.3], *p* = 0.91 for Wilcoxon test, Column 1 of Figure 2A-B). Note that the confidence interval is long, so moderate effects in either direction cannot be ruled out (two animals showed increased anxiety levels and four decreased levels). Pilo treatment for socially housed mice produced a large increase in anxiety scores (*M*_diff_ = 0.19 95% CI[0.12, 0.24], *d* = 3.2 95% CI[1.6, 5.5], *p* = 0.03 for Wilcoxon test, Column 2 of Figure 2A-B).

We then compared the magnitudes of the changes induced by housing and treatment (Figure 2C, left). We found a substantial effect of Pilo treatment: *M*_diff/diff_ = 0.19 95% CI[0.05, 0.30], *d* = 1.6 95% CI[0.4, 4.3], *p* = 0.07 for Mann-Whitney test. Note that there is considerable uncertainty about the magnitude of the Pilo effect under this condition, with the confidence interval stretching down towards negligible effects. This uncertainty is due to high variation in change scores in the standard social housing mice; the overlap in the change scores of SC-alone vs. SC+Pilo pushing up the *p*-value just above the traditional significance level.

We then investigated the effects of social isolation and epilepsy. Consistent with the literature on social deprivation, isolated housing produced a marked increase in anxiety index scores, with all animals increasing from pretest to posttest (*M*_diff_ = 0.11 95% CI[0.06, 0.19], *d* = 1.8 95% CI[1.1, 2.8], *p* = 0.03 for Wilcoxon test, Column 3 of Figure 2A-B). Pilo treatment for isolated mice produced an even larger increase in anxiety scores (*M*_diff_ = 0.33 95% CI[0.27, 0.40], *d* = 5.0 95% CI[4.0, 6.0], *p* = 0.03 for Wilcoxon test, Column 4 of Figure 2A-B). Comparing these changes (Figure 2C, right) shows that Pilo treatment produces a large change above and beyond the effect of isolated housing: *M*_diff/diff_ = 0.21 95% CI[0.13, 0.28], *d* = 3.0 95% CI[1.5, 4.8], *p* = 0.005 for Mann-Whitney test. The confidence interval suggests that all compatible effect sizes are large.

The estimated effects of Pilo treatment were similar for both isolated (*M*_diff/diff_ = 0.21, 95% CI[0.13, 0.28]) and socially housed animals (*M*_diff/diff_ = 0.19, 95% CI[0.05, 0.30]). Thus, the data do not provide strong evidence that housing condition modifies the effect of Pilo treatment, and a formal test for an interaction was not significant (*F*_(1,20)_ = 0.11, *p* = 0.74 for a Treatment x Housing interaction in a non-parametric ANOVA performed on change scores). This means that “epilepsy” increases anxiety levels independently from the housing conditions and that the social isolation-induced increase in anxiety level adds to the “epilepsy”-induced one, resulting in highest anxiety levels in the Pilo-IC group as compared to all other groups.

### Effects of housing and epilepsy in the Novel Object Recognition test

Next, we assessed the effect of social isolation on memory performance using the NOR test. As for anxiety levels, cognitive performance was similar at pretest across groups, with each group mean within 6% of the others (Figure 3A, compare Pre scores across groups). There was notable diversity, though, in pretest scores, with some animals scoring relatively low (50%) and others relatively high (90%). These differences, however, were not clearly predictive of posttest scores (*r* = −0.297 95% CI[-0.626, 0.121], *p* = 0.16).

**Figure 3.**
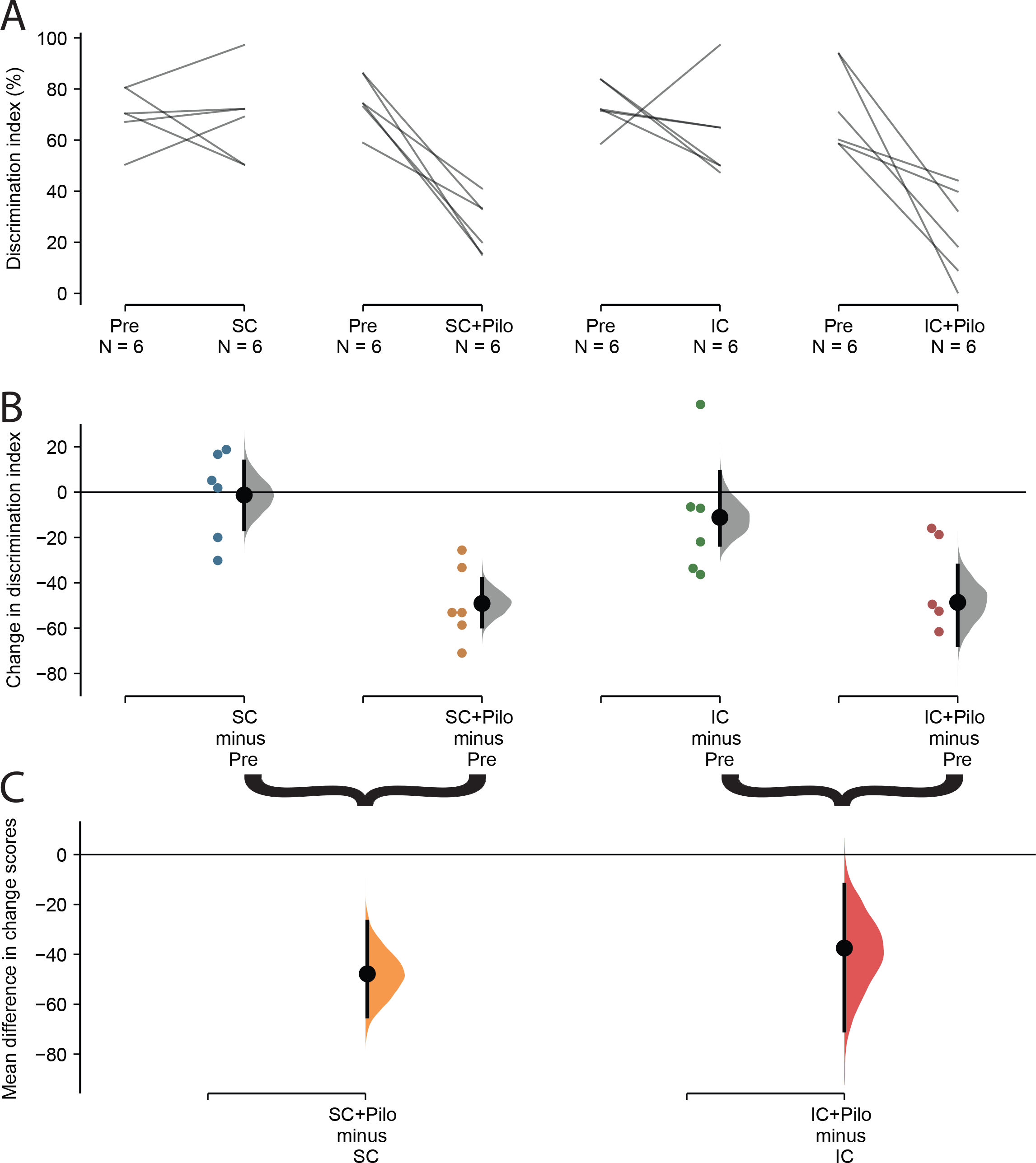
Effect of social isolation on the novel object recognition test in control and epileptic mice. The paired mean difference for four comparisons between posttest (Non-pilo SC, Pilo SC, Non-pilo IC, and Pilo IC) and pretest (Pre) conditions are shown in the Cumming estimation plot. The raw data is plotted on the upper axes; each paired set of observations is connected by a line (**A**). On the lower axis (**B**), each paired mean difference is plotted as a bootstrap sampling distribution. Mean differences are depicted as black dots; 95% confidence intervals are indicated by the ends of the vertical error bars. Colored dots correspond to the posttest-pretest differences for each mouse. As for anxiety levels, although the discrimination index appears similar in average in pretest conditions, we note a wide dispersion of the raw data, i.e. individual animals can have very different levels of performance in the NOR test (compare pretest values in panel A). **(B)** There was no change in performance in the control group (Non-pilo mice maintained in social housing) although individuals displayed variability (first panel from the left). Social isolation in Non-pilo animals (third panel from the left) did not change the performance at the group level (the confidence interval crosses the null line), but we note that five out of six of the isolated animals displayed a decrease in performance as compared to pretest. The changes in discrimination index were substantial in both Pilo-SC (second panel) and Pilo-IC (fourth panel) groups. **(C)** Each mean difference in changes in discrimination index score is plotted as a bootstrap sampling distribution. Mean differences are depicted as dots; 95% confidence intervals are indicated by the ends of the vertical error bars. Epilepsy (in Pilo mice) produced the same increase in anxiety levels in SC and IC conditions.

For animals maintaining social housing during the posttest period, there was no average change in dicrimination index scores (Mean difference score (*M*_diff_) = −1.29% 95% CI[-16.4, 13.5], *d* = −0.08 95% CI[-1.71, 1.05], *p* = 0.92 for Wilcoxon test, Column 1 of Figure 3A-B). Note that the confidence interval is long, so moderate effects in either direction cannot be ruled out (two animals showed similar performance, two increased and two decreased). Pilo treatment for socially housed mice produced a large decrease in performance in the NOR test (*M*_diff_ = −49% 95% CI[-59, −38], *d* = −4.69 95% CI[-6.65, −3.03], *p* = 0.03 for Wilcoxon test, Column 2 of Figure 3A-B).

We then compared the magnitudes of the changes induced by housing and treatment (Figure 3C, left). We found a large effect of Pilo treatment: *M*_diff/diff_ = −47% 95% CI[-65, −27], *d* = −2.6 95% CI[-4.4, −1.3], *p* = 0.008 for Mann-Whitney test.

We then investigated the effects of social isolation and epilepsy. Isolated housing produced a modest decrease in performance (*M*_diff_ = −11% 95% CI [−23, 9], *d* = −0.8 95% CI[− 2.9, 0.8], *p* = 0.3 for Wilcoxon test, Column 3 of Figure 3A-B). Note that there is considerable uncertainty about the magnitude of the isolation effect, with the confidence interval stretching from positive no negative values. This uncertainty is due to one animal, which performance dramatically increased once isolated, pushing up the *p* value above the traditional significance level. If we do not take this animal in consideration, the decrease in performance was large (*M*_diff_ = 21.1% 95.0% CI[ −29.4, −12.3], *d* = −2.75 95.0% CI[-6.11, −2.02], *p* = 0.043). Replication studies are now required to determine whether most control animals perform less once isolated whilst others show an increase of performance (e.g. the animal suffered/was dominated when socially housed). However, we can reasonably propose that social isolation decreased performance in the NOR test is most control mice. Pilo treatment for isolated mice produced a very large decrease in performance (*M*_diff_ = −49% 95% CI[-68, −32], *d* = −2.8 95% CI[-3.7, −2.0], *p* = 0.02 for Wilcoxon test, Column 4 of Figure 3A-B). Comparing these changes (Figure 3C, right) shows that Pilo treatment produces a robust change above and beyond the effect of isolated housing: *M*_diff/diff_ = −38% 95% CI[-71, −12], *d* = −1.3 95% CI[ −2.4, −0.2], *p* = 0.044 for Mann-Whitney test. The confidence interval suggests that all compatible effect sizes are large. The estimated effects of Pilo treatment were similar for both isolated (*M*_diff/diff_ = −38%, 95% CI[-74, −1]) and socially housed animals (*M*_diff/diff_ = −48%, 95% CI[-71, −24]). The large confidence interval for isolated animals is due to the Non-pilo animal showing increased performance after isolation. Thus, the data do not provide strong evidence that housing condition modifies the effect of Pilo treatment, and a formal test for an interaction was not significant (*F*_(1,20)_ = 0.28, *p* = 0.6 for a Treatment x Housing interaction in a non-parametric ANOVA performed on change scores). This means that “epilepsy” decreases cognitive performance in the NOR test independently from the housing conditions, but that the social isolation-induced decrease in performance adds to the “epilepsy”-induced one, resulting in poor performance in the Pilo-IC group as compared to all other groups.

We conclude that housing conditions affect cognitive performance in the NOR test in most individuals (although not for the group) and that epilepsy is associated with similar substantial cognitive deficits in the NOR test irrespective of housing conditions.

### Anxiety and novel object recognition performance co-vary independently of housing and treatment conditions

Anxiety correlates with poor performance in a variety of cognitive tasks (Moran, 2016). Given the wide distribution of data points for the two behavioral tests used here, we tested for a correlation between the two variables. We found a strong linear relationship between anxiety levels and the discrimination index, *r* = −0.858 95% CI[-0.918, −0.758], *p* < 0.0001, (Figure 4A). In pretest, all animals were distributed around the regression line, spanning various levels of anxiety/discrimination values, reflecting the individual differences between mice in pretest condition. In posttest conditions, mice that remained socially housed were distributed within the pretest values. Social isolation in the control (Non-pilo) group did not change the anxiety/discrimination relationship but shifted the points towards higher anxiety/lower performance. Pilo treatment in socially housed mice resulted in a further shift while combining both Pilo treatment and isolation produced the maximum shift. Thus, the relationship between anxiety/discrimination values remained constant independently of the experimental conditions, i.e. there is a strong and stable rule of covariance between anxiety and discrimination levels. The experimental conditions appear only to change the position along the regression line.

**Figure 4:**
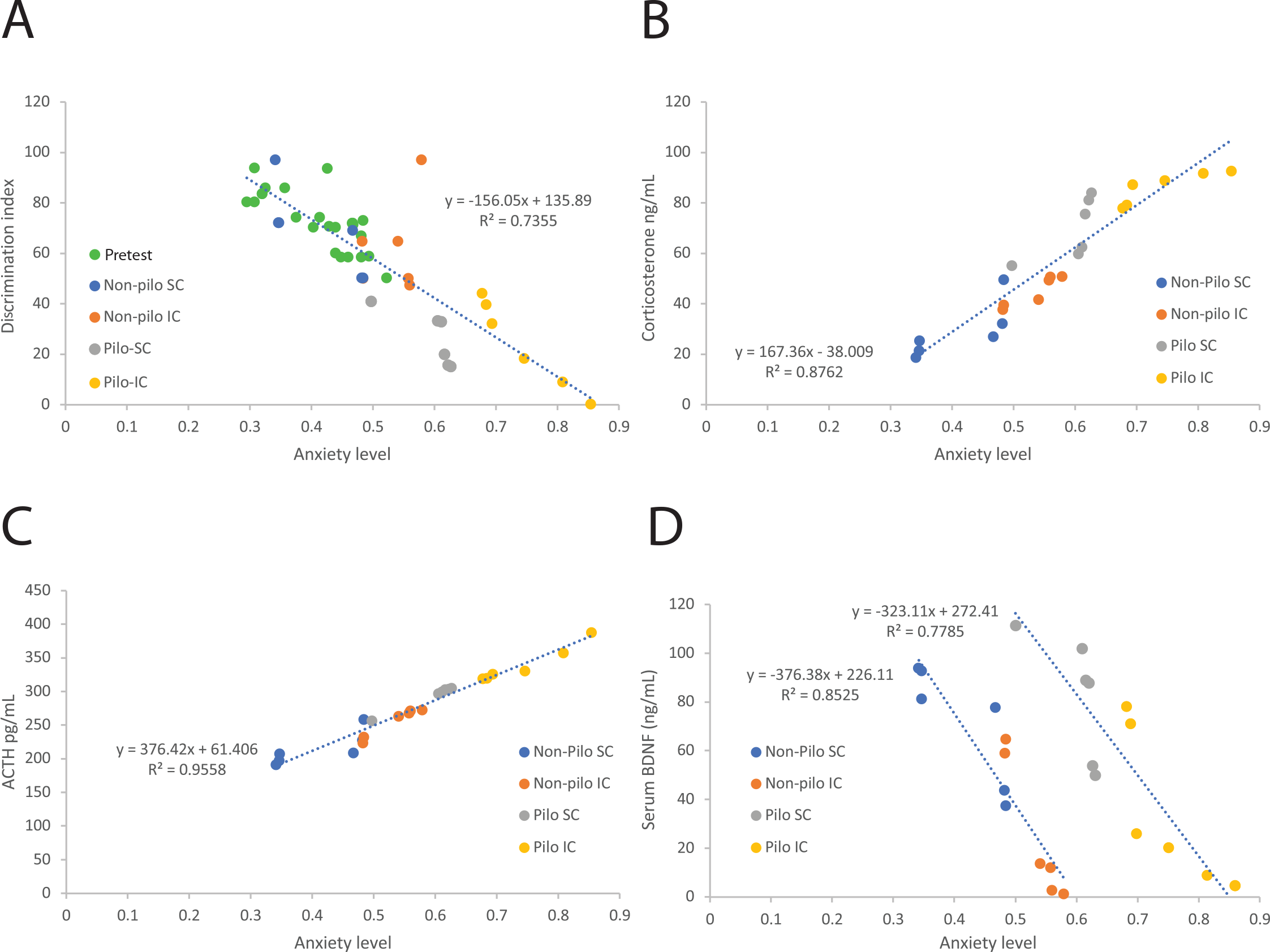
Co-variance between anxiety levels and the discrimination index, corticosterone, ACTH and BDNF levels. **(A)** The discrimination index is linearly correlated to anxiety levels. All mice are displayed. The dispersion of pairs of values for anxiety levels (Figure 2A) across 0.25 unit and discrimination index (Figure 2B) across 50 units at baseline is clearly apparent (green dots). All mice are distributed close to the regression line (built from all data points). The Non-pilo group maintaining social housing (dark blue dots) remained in the cloud of baseline values. The Non-pilo isolated group (orange dots) showed a downward shift. Note the presence of the outlier with high cognitive performance and high anxiety level. Pilo animals displayed a further downward shift with higher anxiety levels and lower cognitive performance, with isolated animals showing the strongest phenotype. Corticosterone **(B)** and ACTH **(C)** levels are strongly linearly correlated to anxiety levels. As for the discrimination index, the phenotype increased in a group-dependent manner Non-pilo SC < Non-pilo IC < Pilo SC < Pilo IC. **(D)** The relationship between BDNF and anxiety levels was also linear but treatment-dependent. The regression line was shifted to the right (greater anxiety levels) in epileptic mice.

### Social isolation and epilepsy increase corticosterone and ACTH levels

We then assessed biological markers obtained at the end of the experimental protocol (Figure 1A). Using treatment as reference (Figure 5A, right panel), the unpaired mean difference between Pilo-SC and Non-pilo-SC corticosterone levels was 40.7 ng/mL [95.0%CI 27.2, 51.5] and the unpaired mean difference between Pilo-IC and Non-pilo-IC was 41.3 ng/mL [95.0%CI 34.6, 47.2], corresponding to an increase of corticosterone by 55% and 91%, respectively. In both conditions, the two-sided *P* value of the Mann-Whitney test was 0.00507. The unbiased Cohen’s *d* was 3.228 and 6.204 for the SC and IC groups, respectively. The difference of differences (Figure 6A) was 0.595 ng/mL [95.0%CI −15.202, 16.392], indicating no significant interaction between treatment and housing. Using social housing as reference (Figure 5A, left panel), the unpaired mean difference between Non-pilo-SC and Non-pilo-IC corticosterone levels was −16.0 ng/mL [95.0%CI −23.5, −4.43] and the unpaired mean difference between Pilo-SC and Pilo-IC corticosterone levels was −16.6 ng/mL [95.0%CI −26.4, −6.68], corresponding to an increase in corticosterone level in the isolated group by 59% and 95%, respectively. In both conditions, the two-sided *P* value of the Mann-Whitney test was 0.0306. The unbiased Cohen’s *d* was 1.656 and 1.591 for the Non-pilo and Pilo groups, respectively. The effect of isolation had similar effects in both control and pilo groups (1.6 standard deviation). We conclude that both epilepsy and social isolation produce a significant increase in corticosterone levels, the most substantial effect occurring in the Pilo-IC group.

**Figure 5:**
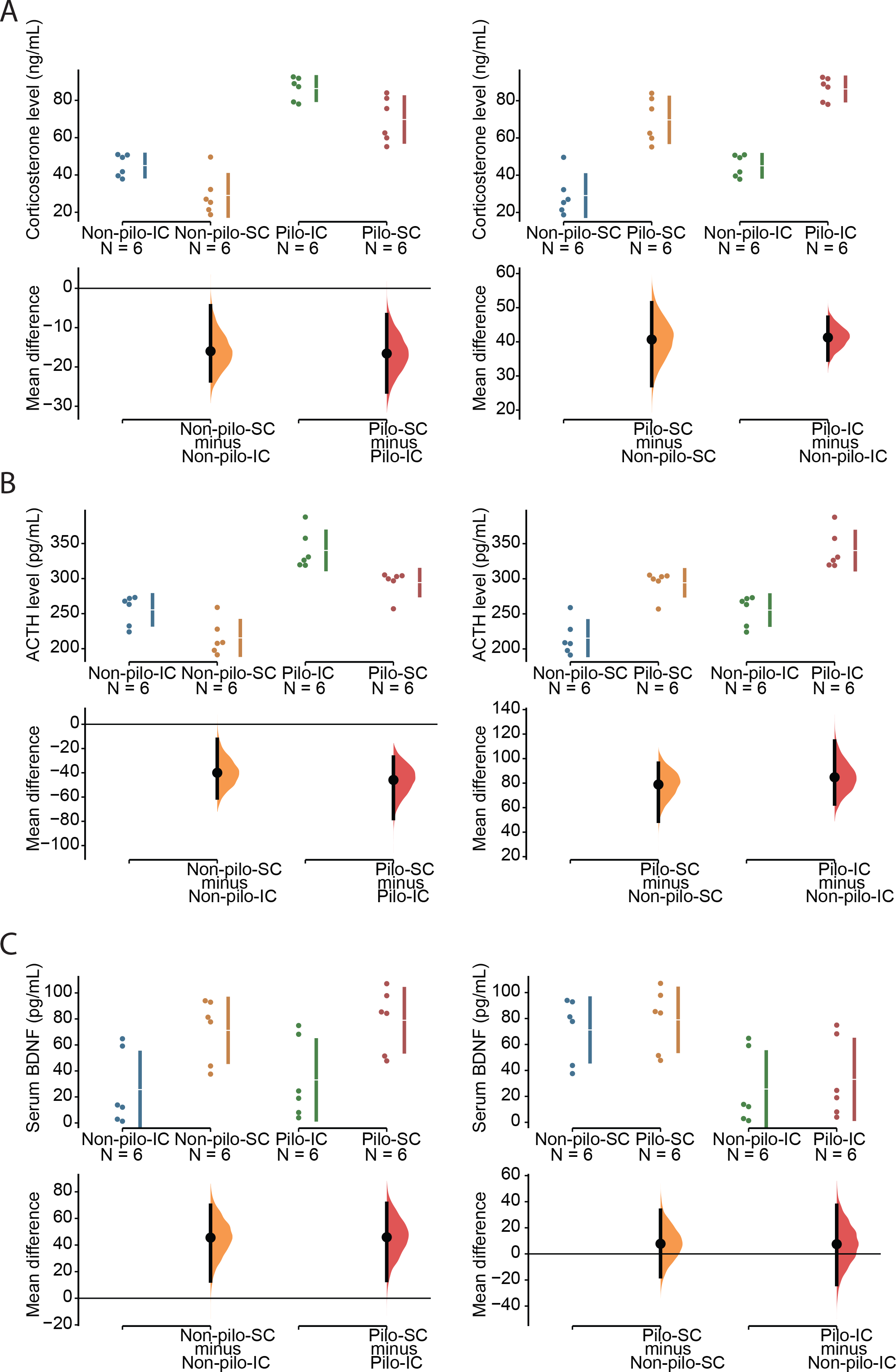
Effect of social isolation on corticosterone, ACTH and BDNF levels in control and epileptic mice. In panels A, B and C, the mean difference for two comparisons is shown in the Cumming estimation plot. All raw data is shown on the upper axes. For each group, summary measurements (mean ± standard deviation) are shown as gapped lines. Each mean difference is plotted on the lower axes as a bootstrap sampling distribution. Mean differences are depicted as black dots; 95% confidence intervals (CIs) are indicated by the ends of the vertical error bars. **(A)** Social isolation increased corticosterone levels by 15 units in both Non-pilo and Pilo groups (left panel). The effect of epilepsy was even stronger (40 units) in the SC and IC groups (right panel). **(B)** Social isolation increased ACTH levels by 40 units in both Non-pilo and Pilo groups (left panel). The effect of epilepsy was even stronger (80 units) in the SC and IC groups (right panel). **(C)** Social isolation decreased BDNF levels by 40 units in both Non-pilo and Pilo groups (left panel). The Epilepsy condition had no apparent effect on BDNF levels in both SC and IC groups.

**Figure 6:**
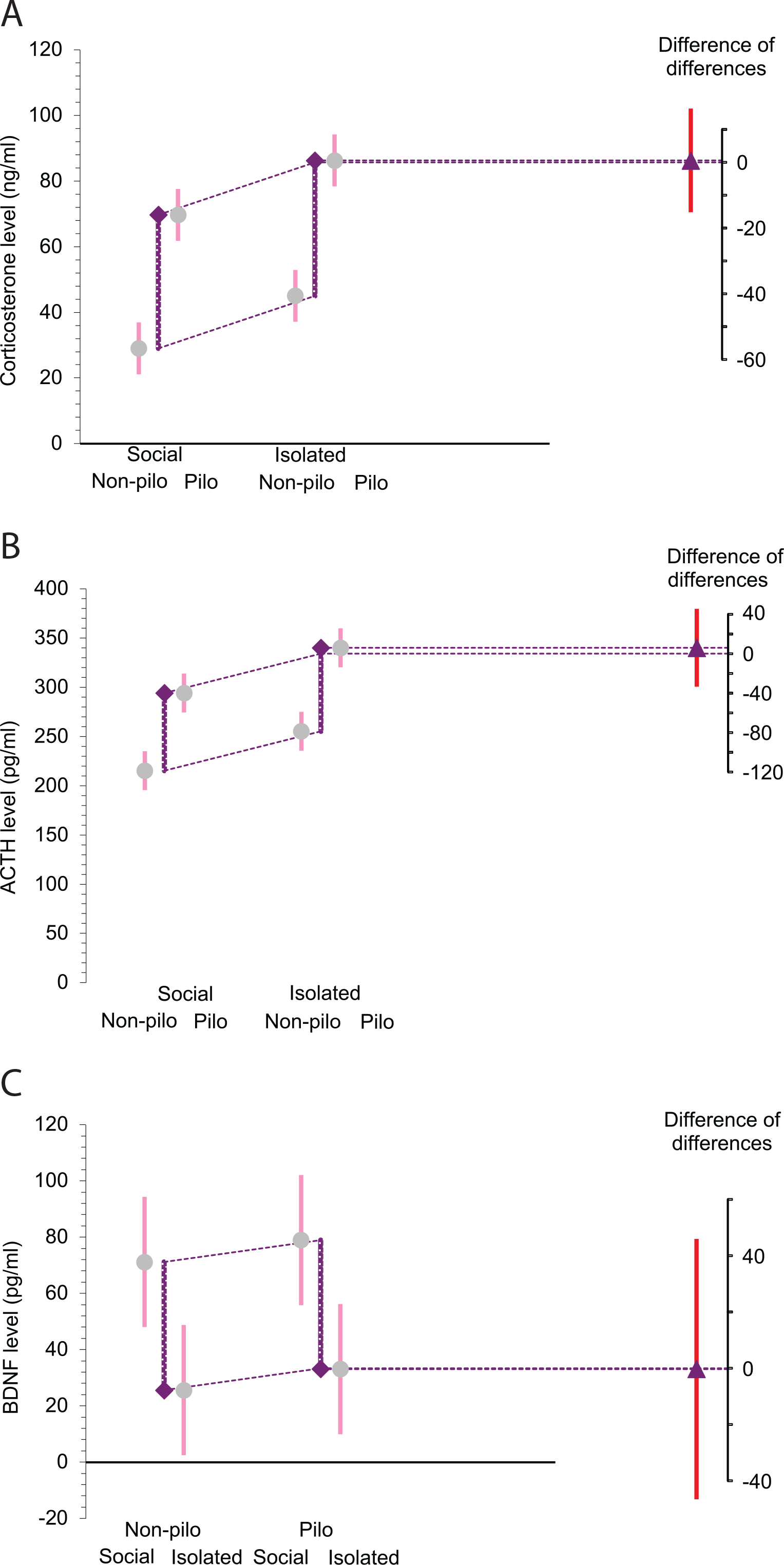
Housing x treatment interaction on corticosterone, ACTH and BDNF levels. Means and 95% CIs are as in Figure 5. The (Pilo – Non-pilo) differences between the right two means and the left two means are marked by the purple lines ending in diamonds (indicating the direction of the change). The slanted dotted lines make a comparison of the two differences. The difference between the differences is marked by the triangle on the difference axis, with its CI. The analysis of the difference of differences did not reveal apparent housing x treatment interaction for corticosterone **(A)**, ACTH **(B)** and BDNF **(C)** levels.

For ACTH, using Non-pilo treatment as reference (Figure 5B, right panel), the unpaired mean difference between Pilo-SC and Non-pilo-SC ACTH levels was 78.9 pg/mL [95.0%CI 48.9, 96.3] and the unpaired mean difference between Pilo-IC and Non-pilo-IC was 84.8 pg/mL [95.0%CI 63.0, 1.14e+02], corresponding to an increase in ACTH level in the Pilo group by 37% and 33%, respectively. The two-sided *P* values of the Mann-Whitney test were 0.00824 and 0.00507, respectively. The unbiased Cohen’s *d* was 3.336 and 3.189 for the SC and IC groups, respectively, suggesting that epilepsy had strong but similar effects (3 standard deviation) in both socially housed and isolated mice. The difference of differences (Figure 6B) was 5.914 pg/mL [95.0%CI −33.628, 45.457], indicating no significant interaction between treatment and housing. Using social housing as reference (Figure 5B, left panel), the unpaired mean difference between Non-pilo-SC and Non-pilo-IC ACTH levels was −40.0 pg/mL [95.0%CI −60.8, −12.4] and the unpaired mean difference between Pilo-SC and Pilo-IC ACTH levels was −45.9 pg/mL [95.0%CI −77.7, −27.0], corresponding to an increase in ACTH level in the isolated group by 21% and 17%, respectively. The two-sided *P* values of the Mann-Whitney test were 0.0202 and 0.00507, respectively. The unbiased Cohen’s *d* was 1.596 and 1.817 for the Non-pilo and Pilo groups, respectively. The effect of isolation had similar effects in both control and Pilo groups (1.6 standard deviation). We conclude that both epilepsy and social isolation produce a significant increase in ACTH levels, the largest effect occurring in the Pilo-IC group.

Since ACTH and corticosterone levels are stress markers, not surprisingly, we found a strong linear relationship between anxiety levels and corticosterone *r* = 0.936 95% CI[0.856, 0.972], *p* < 0.0001 (Figure 4B) as well as for ACTH *r* = 0.978 95% CI[0.948, 0.99], *p* < 0.0001 (Figure 4C) levels. We note again the strong co-variance of anxiety level with corticosterone/ACTH levels. The distribution of points along the regression line reflects the variability of individuals across housing/treatment conditions. The housing/treatment conditions do not impact upon the co-variance rule itself. As for the discrimination index (Figure 4A), the conditions merely shift the distribution of points toward higher values as a function of the severity of the effect, the Pilo isolated group displaying the greater anxiety-corticosterone-ACTH levels, and the control Non-pilo social the lowest levels.

### Social isolation decreases serum BDNF levels in control and pilo mice

Strong stressful situations can trigger a state of vulnerability to depression in some individuals, which can be identified with low serum BDNF levels (Blugeot et al., 2011; Becker et al., 2015). We thus assessed serum BDNF in the different groups. Using Non-pilo treatment as reference (Figure 5C, right panel), the unpaired mean difference between Pilo-SC and Non-pilo-SC BDNF levels was 7.76 pg/mL [95.0%CI −17.5, 33.5] and the unpaired mean difference between Pilo-IC and Non-pilo-IC was 7.45 pg/mL [95.0%CI −23.5, 37.3]. The two-sided *P* values of the Mann-Whitney test were 0.378. The unbiased Cohen’s *d* was 0.293 and 0.232 for the SC and IC groups, respectively. These results suggest that epilepsy does not affect significantly serum BDNF levels in mice in both housing conditions. Using social housing as reference (Figure 5C, left panel), the unpaired mean difference between Non-pilo-SC and Non-pilo-IC BDNF levels was 45.5 pg/mL [95.0%CI 13.0, 69.9] and the unpaired mean difference between Pilo-SC and Pilo-IC was 45.9 pg/mL [95.0%CI 13.4, 71.3], corresponding to a decrease in the isolated group by 57% and 51%, respectively. The two-sided *P* values of the Mann-Whitney test were 0.0306 for Non-pilo and Pilo groups. The unbiased Cohen’s *d* was −1.577 and −1.527 for the Non-pilo and Pilo groups, respectively. The effect of isolation had similar effects in both control and Pilo groups (1.5 standard deviation). The difference of differences (Figure 6C) was −0.312 pg/mL [95.0%CI −46.608, 45.983], indicating no significant interaction between treatment and housing. Thus, housing conditions has strong effects on serum BDNF levels in both Pilo and Non-pilo groups.

In contrast with the condition-independent co-variance rules found previously for corticosterone and ACTH, individual BDNF data show a more complex relationship. In both control (Non-pilo) and epilepsy (Pilo) conditions, we found a strong linear relationship between anxiety levels and serum BDNF levels, the housing condition just shifting the distribution of points, *r* = −0.923 95% CI[-0.948, −0.7439], *p* < 0.0001 and *r* = −0.882 95% CI[-0.967, −0.625], *p* < 0.0001 for Non-pilo and Pilo groups, respectively (Figure 6D). However, the epilepsy condition clearly shifts the relationship to the right of the graph, i.e. as compared to the control condition, a similar level of serum BDNF level is associated with a higher level of anxiety in Pilo mice. Said differently, the epilepsy condition changes the setpoint (the *y* intercept of the linear regression function) of the relationship between anxiety and serum BDNF levels, without an apparent change of the slope (the co-variance rule).

Together, the results obtained in control (Non-pilo) mice demonstrate that social isolation has a strong effect on many behavioral and biological stress indicators. They recapitulate the main features of social isolation performed after weaning to model early life stress (Mumtaz et al., 2018). In animals maintained in groups, epilepsy had a direct effect on stress indicators. These effects were exacerbated in singly housed Pilo mice. We conclude that social isolation produces a stronger phenotype (for the tests used here) in the Pilo mouse experimental model of epilepsy. The mouse study was designed to evaluate two independent variables (housing and epilepsy) on several parameters relevant to stress and cognitive performance. A limitation of this study is that we did not evaluate spontaneous seizures with video/EEG. For this, we would have had to double the number of animals to evaluate the two independent variables. Indeed, the surgery, the implantation of an invasive device, and the treatment (anesthesia, anti-inflammatory) may by themselves change the animal’s biology and thus change the relationships between anxiety and cognitive performance, etc. requiring a study design including animals equipped with EEG transmitters and not-equipped animals. Future studies should address this issue. However, we directly investigated the effect of social conditions on seizures in the pilocarpine rat model.

### Effect of social isolation in control rats

Since we could not perform a full two-condition study like in mice, we first wanted to determine whether the general effects of isolation in control animals were similar in the two species, simplifying the experimental protocol in control rats (Figure 7A). We compared three groups of rats: singly housed animals (Isolated), animals kept in pairs (Paired) and isolated animals interacting with the experimentor every day (Handled). The rationale for designing these three groups is the constraints imposed by the experimental part done in the pilocarpine model of epilpsy (see below). Because of their larger size, it was not technically possible to maintain rats in small colonies while performing wireless 24/7 recordings. For the social group, animals were kept in pairs. The justification of the Handled group is that some EEG studies are performed with wired systems, which precludes group housing, as animals tend to sever the wires.

**Figure 7:**
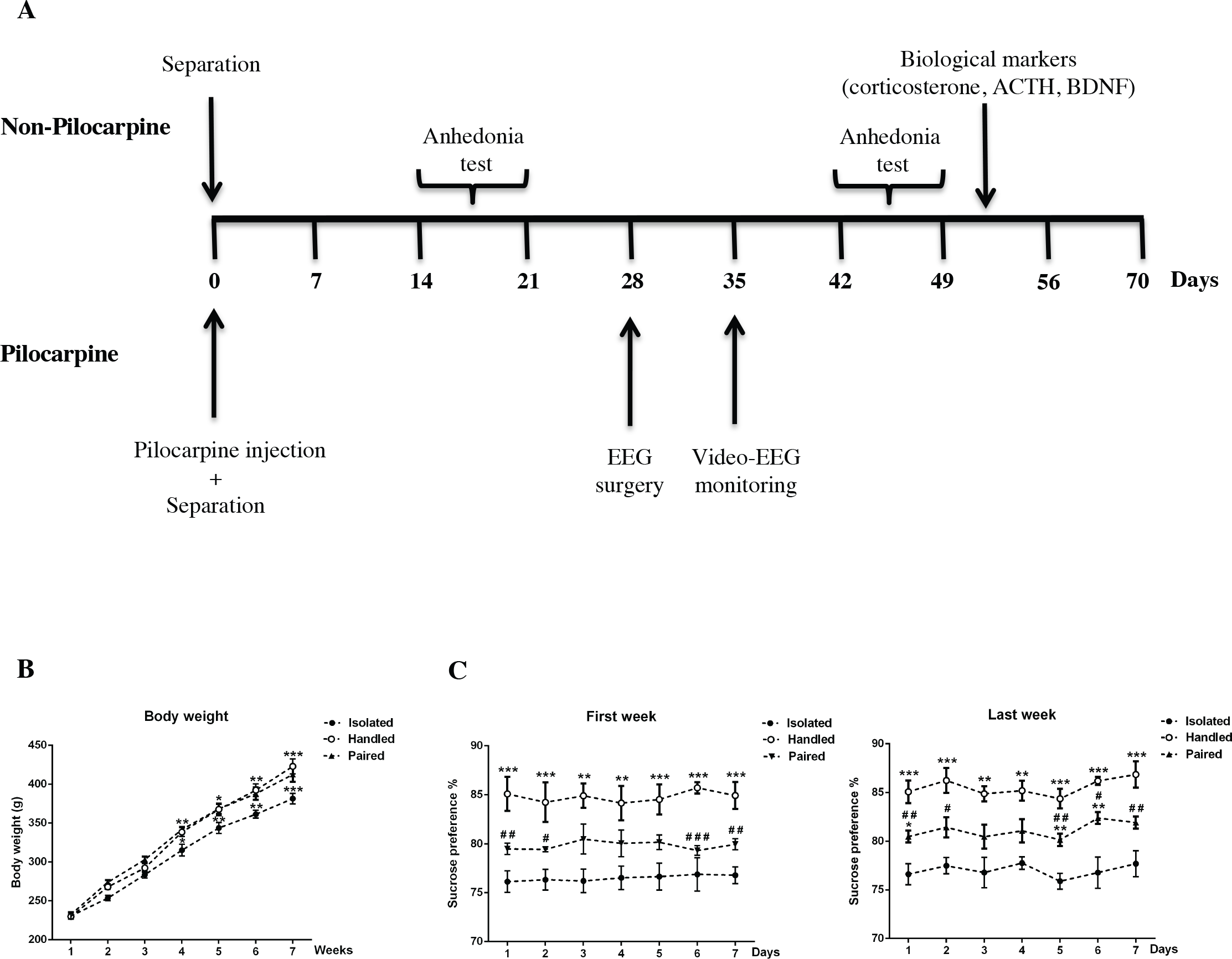
Effect of social isolation on body weight and anhedonia in control rats. **(A)** Experimental protocol in Non-pilo and Pilo rat groups. In the control (Non-pilo) group (top part of the time axis) rats were assigned at Day 0 to Isolated, Handled and Paired groups. Anhedonia was assessed two weeks and six weeks later. During week 7, animals were killed for the analysis of corticosterone, ACTH and BDNF levels. In the Pilo group (bottom part of the time axis), animals were injected with pilocarpine at Day 0 and assigned to Isolated, Handled and Paired groups. Four weeks later, rats were equipped with wireless EEG transmitters, and recordings started one week later for three weeks. **(B)** Evolution of the average body weight over seven weeks in control (Non-pilo) rats for the three Isolated, Handled and Paired groups. The isolated group significantly gained less weight. (**C**) In control rats, Isolated animals already showed anhedonia during the first evaluation (two weeks after separation) as compared to Paired animals. Interestingly, Handled animals showed increased sweet water consumption as compared to the Paired group. The same result was found during the second evaluation period six weeks after group assignment. Data are mean ± SEM. \**p*<0.05, \*\**p*<0.01 and \*\*\**p*<0.001 in comparison with the isolated group; #*p*<0.05, ##*p*<0.01 and ###*p*<0.001 in comparison with the handled group.

We first tested the effect of isolation in control rats. The ANOVA repeated measures showed a significant effect of social condition (F_(2,17)_=41.58, *p*<0.0001d) and time (F_(6,102)_ =266.83, *p*<0.0001d) on the gain of body weight (Figure 7B). At the 4^th^ week, *post-hoc* analysis showed that the Isolated group gained significantly less weight (by 7%) than the Handled and the Paired group. There was no significant difference between all groups from the 1^st^ to the 3^rd^ week.

We used the sucrose preference test to assess anhedonia. ANOVA repeated measures showed a main effect of social isolation on sweet water consumption (Figure 7C) already during the third week (F_(2,12)_=21.66, *p*<0.0001^e^) up to the last week (F_(2,12)_=97.85, *p*<0.0001^f^) with no time effect in both weeks (first week: F_(6,72)_ =0.11, *p*>0.05 and last week: F_(6,72)_ = 1.09, *p*>0.05; respectively ^e,f^). During the third and the last week, *the post-hoc* analysis revealed that sweet water consumption was significantly lower (by 5-10%) in the Isolated than in the Handled (*p*<0.01, *p*<0.001) and Paired groups (*p*<0.05, *p*<0.01 and *p*<0.001). Thus, just two weeks of social isolation is sufficient to produce a state of anhedonia in control rats.

We then tested stress-related corticosterone and ACTH hormones in control rats. For corticosterone, using the isolated condition as reference (Figure 8A), the unpaired mean difference between Isolated and Handled corticosterone levels was −8.17 ng/mL [95.0%CI −12.2, −4.2] and the unpaired mean difference between Isolated and Paired was −7.04 ng/mL [95.0%CI −11.1, −3.47], corresponding to a 60% and 48% increase in Isolated animals, respectively. The two-sided P values of the Mann-Whitney test were 0.0216 and 0.0167, respectively. The unbiased Cohen’s *d* was −2.095 and −1.926 for the Handled and Paired groups, respectively. Social isolation thus had a strong effect on corticosterone levels in control rats. The two distributions for the Handled and Paired groups were similar (unbiased Cohen’s *d* was 0.367). For ACTH, using the isolated condition as reference (Figure 8B), the unpaired mean difference between Isolated and Handled corticosterone levels was −1.14e+02 pg/mL [95.0%CI −2.22e+02, −63.7] and the unpaired mean difference between Isolated and Paired was −1.07e+02 pg/mL [95.0%CI −1.48e+02, −50.5], corresponding to a 52% and 47% increase in Isolated animals, respectively. The two-sided P values of the Mann-Whitney test were 0.0122 and 0.0235, respectively. The unbiased Cohen’s *d* was −1.579 and −1.685 for the Handled and Paired groups, respectively. Social isolation thus had also a strong effect on ACTH levels. The two distributions for the Handled and Paired groups were similar (unbiased Cohen’s *d* was 0.086). Finally, we assessed serum BDNF levels. Using the isolated condition as reference (Figure 8C), the unpaired mean difference between Isolated and Handled BDNF levels was 0.32 pg/mL [95.0%CI 0.151, 0.524] and the unpaired mean difference between Isolated and Paired was 0.458 pg/mL [95.0%CI 0.198, 0.697], corresponding to a 16% and 22% decrease in Isolated animals, respectively. The two-sided P values of the Mann-Whitney test were 0.0216 and 0.012, respectively. The unbiased Cohen’s *d* was 1.749 and 1.464 for the Handled and Paired groups, respectively. Social isolation thus had also a strong effect decreasing BDNF levels in rats, as in control mice (Figure 5). The two distributions for the handled and paired groups were similar (unbiased Cohen’s *d* was 0.457).

**Figure 8:**
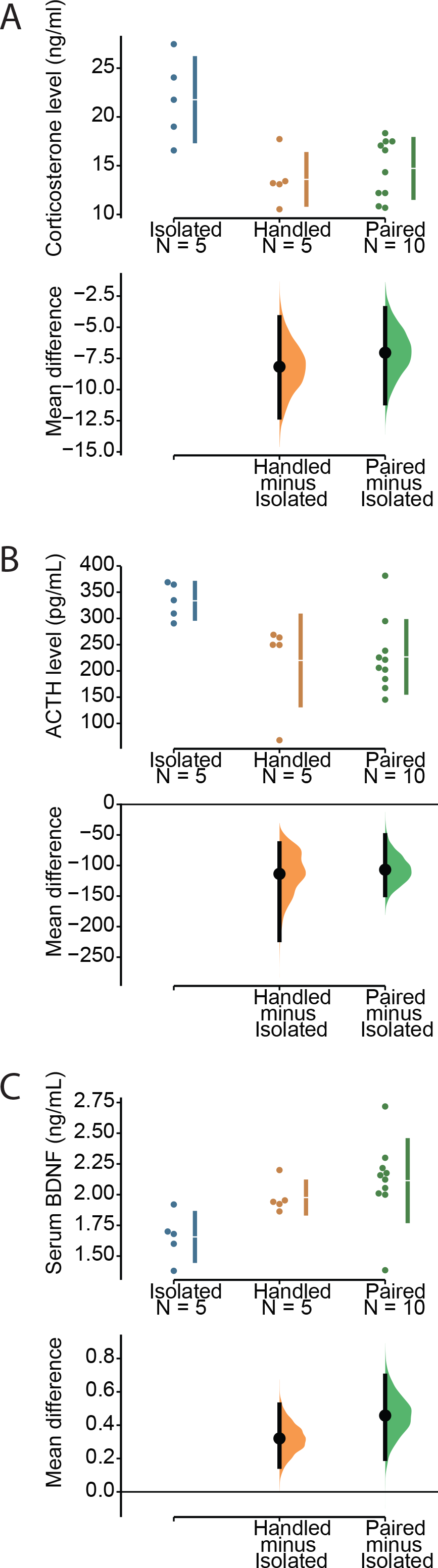
Effect of social isolation on corticosterone, ACTH and BDNF levels in control rats. **(A)** Social isolation increased corticosterone levels by seven units in Isolated rats as compared to Handled and Paired animals. **(B)** Social isolation increased ACTH levels by 100 units in Isolated rats as compared to Handled and Paired animals. **(C)** Social isolation decreased BDNF levels by 0.4 units in Isolated rats as compared to Handled and Paired animals. The effect of isolation was significant for the three biomarkers, the CIs being far from the null value. The overlap of the distributions between Handled and Paired animals suggests a lack of difference between the two conditions.

We conclude that, as in mice, social isolation in control rats produces a phenotype associated with stress. Interestingly, daily handling of otherwise isolated animals prevented the occurrence of the phenotypic markers of stress. They did not defer from paired animals for the measures considered here. We then compared how different housing conditions influence the epilepsy phenotype (the experimental protocol is shown in Figure 7A).

### Social isolation worsens epilepsy severity in Pilo rats

Typical seizures included low voltage fast activity, a tonic phase and a tonic-clonic phase, which were associated with the typical behavioral expression (using video recordings performed during the light phase). Most seizures terminated with a logarithmic slowing down of the activity (based on visual inspection), characteristic of a homoclinic bifurcation (El Houssaini et al., 2015; Saggio et al., 2017). Since the recording device provides an AC signal, we could not detect the eventual presence of DC shift at seizure onset, which would have indicated seizures of the Epileptor class (saddle node/homoclinic), the most common form of seizure dynamics found across species, including in humans (Jirsa et al., 2014). Interictal spikes could precede seizure onset, but this was not found systematically, including within the same animal. We did not find noticeable differences in the EEG between seizures occurring in the three different groups (Isolated, Handled, Paired). The pilocarpine experimental model of epilepsy is characterized by the presence of different types of bursts of spikes, which dynamics are very different from seizures (Chauviere et al., 2012). These bursts of spikes were not considered as seizures.

Since all Pilo rats were EEG recorded 24/7, we first verified the stability of seizure activity per week. Figure 9 shows that the average seizure frequency per week was remarkably stable for each recorded animal in each group, i.e. seizure frequency did not tend to increase or decrease over time. We thus pooled the seizures recorded in each animal over the three weeks to assess the effects of social isolation.

**Figure 9:**
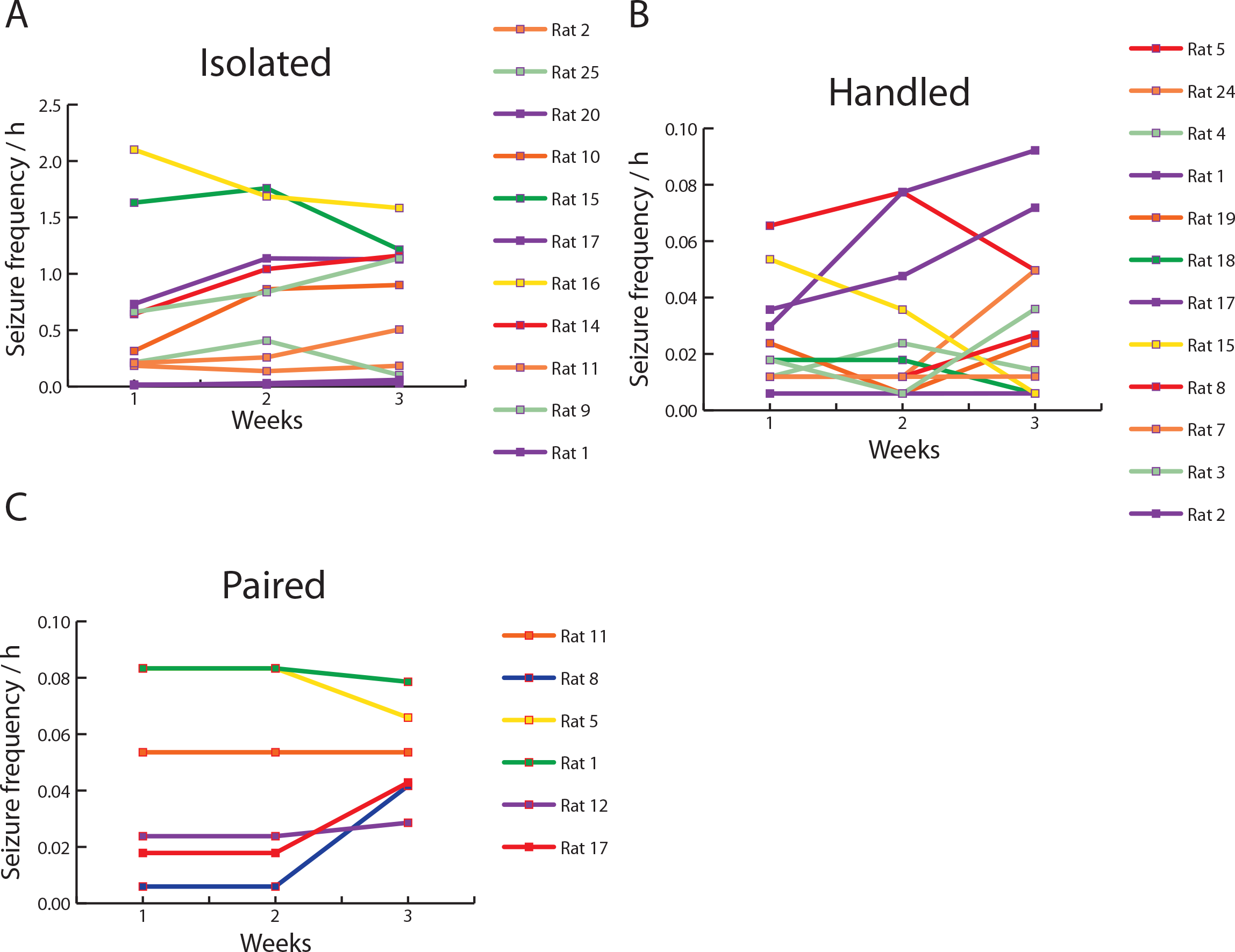
Distribution of seizures per week for each animal. The average seizure frequency (seizures per hour) for each recording week is shown for **(A)** Isolated (n=11), **(B)** Handled (n=12) and **(C)** Paired rats (n=6). Note that the scale of seizure frequency in the Isolated group is one order of magnitude higher than the other two groups. Two animals (out of 11) in the Isolated group had a seizure frequency similar to that found in Paired or Handled animals (0.03/h). The three groups displayed stability (despite some individual variability) over the three weeks of recordings.

The unpaired mean difference in seizure frequency between Isolated and Handled animals (Figure 10A) was −0.665 seizure/h [95.0%CI −1.04, −0.372] and the unpaired mean difference between Isolated and Paired was −0.65 seizure/h [95.0%CI −1.04, −0.356]. The two-sided P value of the Mann-Whitney test were 0.0004 and 0.0104, respectively. The unbiased Cohen’s *d* was −1.547 and −1.26 for the Handled and Paired groups with respect to Isolated, respectively.

**Figure 10:**
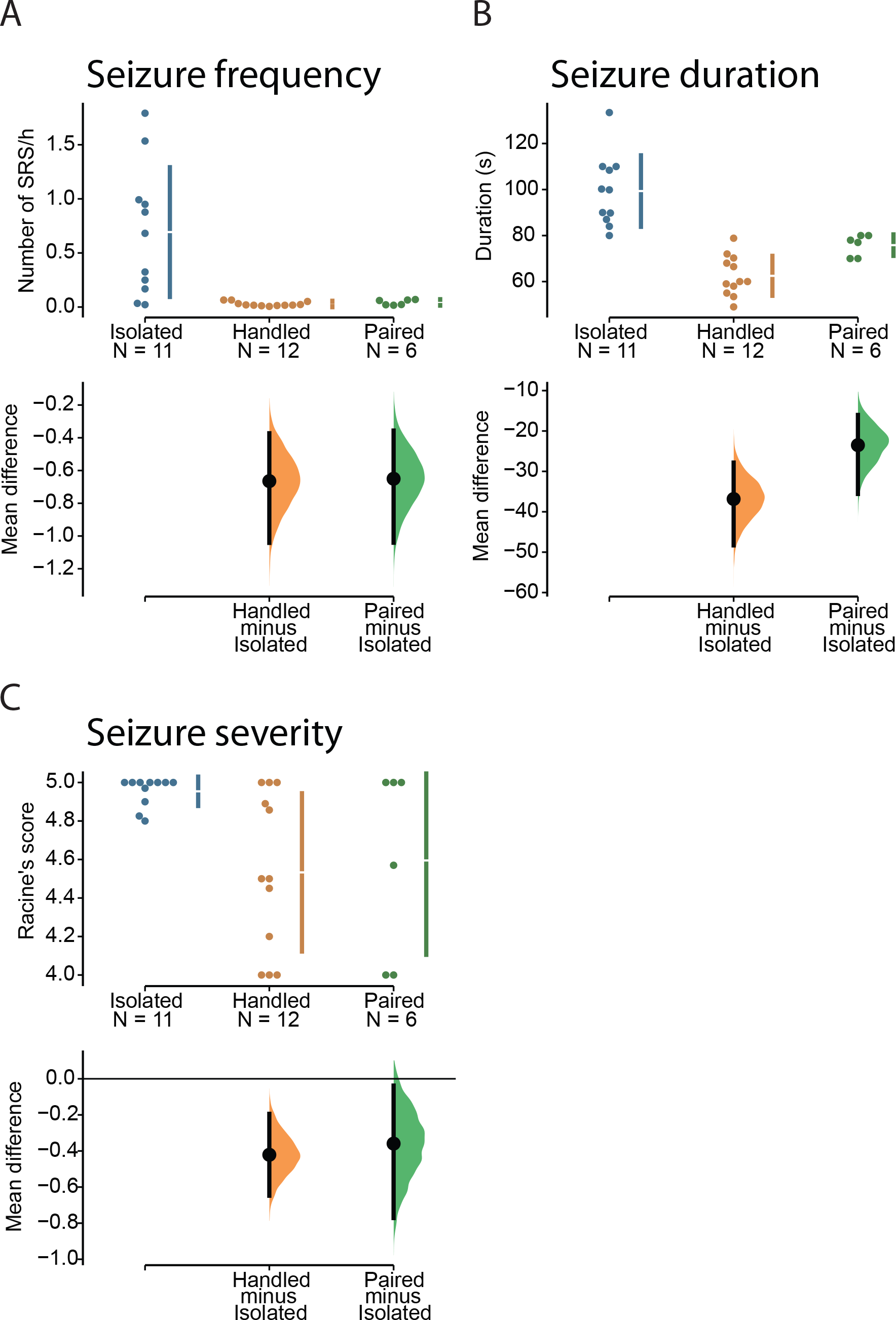
Social isolation increases spontaneous seizures by a factor of 16 in rats. **(A)** The seizure frequency (number of spontaneous recurrent seizures - SRS per hour), **(B)** duration and **(C)** severity are shown for Isolated, Handled and Paired rats (n=11, n=12, and n=6). Isolated rats displayed a very severe epileptic phenotype as compared to the other groups. Handled and Paired groups showed a similar phenotype.

The distributions between Handled and Paired groups were similar (unbiased Cohen’s *d* was 0.623).

The unpaired mean difference in seizure duration between Isolated and Handled (Figure 10B) was −36.8 s [95.0%CI −48.3, −27.8]. The unpaired mean difference between Isolated and Paired was −23.5 s [95.0%CI −35.6, −16.0]. The two-sided P value of the Mann-Whitney test were 5.5e-05 and 0.00149, respectively. The unbiased Cohen’s *d* was −2.857 and −1.72 for the Handled and Paired groups with respect to Isolated, respectively. The distributions between Handled and Paired groups were different (unbiased Cohen’s *d* was 1.658). Although their confidence intervals overlapped (no significant difference in seizure duration), the distribution was narrower in the Paired group overlapping with the distribution of more prolonged duration seizures in the Handled group (Figure 10B).

Seizure severity was assessed with the video during the light phase only. The unpaired mean difference between Isolated and Handled groups (Figure 10C) was −0.421 [95.0%CI - 0.648, −0.194]. The unpaired mean difference between Isolated and Paired was −0.359 [95.0%CI −0.772, −0.0377]. The two-sided P value of the Mann-Whitney test were 0.0121 and 0.26, respectively. The unbiased Cohen’s *d* was −1.343 and −1.177 for the Handled and Paired groups with respect to Isolated, respectively. The distributions between Handled and Paired groups were not different (unbiased Cohen’s *d* was 0.135).

We conclude that social isolation dramatically worsened the seizure phenotype, with seizures being 16 times more frequent in isolated animals as compared to those socially housed or those having daily interaction with experimenters. Interestingly, daily interactions in otherwise singly housed animals had a similar effect on epilepsy severity as compared to animals maintaining social interaction.

## DISCUSSION

Our results show that single housing exacerbates the phenotype in male Wistar rats and Swiss mice in the pilocarpine experimental model of epilepsy. It is important to note that daily social interaction with the experimenter was sufficient to prevent the development of a severe epilepsy phenotype in rats although they were singly housed. It was as efficient as keeping animals in pairs. Although we could not quantify it, we noted that isolated Pilo animals were very aggressive and displayed escaping behavior when cages were changed every week. Their cages were close to one another, suggesting that visual contact was not sufficient to prevent the effect of social isolation. Abnormal reaction to handling is a direct consequence of single housing (Hatch et al., 1965).

In contrast, rats and mice socially housed (or in daily contact with an experimenter) remained calm when handled. In the laboratory, we are also performing electrophysiological recordings with high-density silicon probes in experimental rat models of epilepsy. The equipment is protected with a copper mesh hat, which precludes social housing. We are using large cages, divided in two, separated by a grid allowing physical interaction. Each compartment contains one animal with epilepsy. Animals can make nose contact and interact with their whiskers through the bars. Animals do not show any sign of anxiety, and wire connectors are easily inserted, as in control animals (no struggle). This type of housing allows wired recordings in rats and mice with epilepsy, while maintaining a certain level of social interaction.

Our data on control animals confirms the vast body of literature reporting that social isolation constitutes a major stressful situation that can lead to a depression-like profile (Mumtaz et al., 2018). Among the numerous markers that can be assessed, we focused on general ones such as ACTH, corticosterone, and anhedonia. When rats and mice are singly housed, ACTH and corticosterone levels are increased (Veenema et al., 2005; Djordjevic et al., 2012). Visual inspection of the raw data seems to indicate a bimodal distribution of anxiety levels in pretest conditions in control mice (Figure 2A). It will be interesting to determine whether these animals correspond to animals susceptible and resilient to depression (Han and Nestler, 2017). When exposed to an intense stress (test), control rats also split into two populations, vulnerable and non-vulnerable to depression (Blugeot et al., 2011; Becker et al., 2015; Claverie et al., 2016; Bouvier et al., 2017; Becker et al., 2019). Thus, in pretest conditions, rats and mice are not homogeneous, as they are going to react differently when challenged by external factors such as stress or epileptogenesis (Bernard, 2016; Medel-Matus et al., 2017).

In pilocarpine experimental models of epilepsy, ACTH and corticosterone levels are increased (Mazarati et al., 2009; Inostroza et al., 2012; Ngoupaye et al., 2013). A direct comparison of ACTH and corticosterone values between studies is difficult as they depend upon the time of collection and the way blood is collected (*in vivo* or *post mortem* as done here). Despite the differences in experimental procedures, all results are qualitatively similar (increase due to social isolation or epilepsy). To the best of our knowledge, we here provide the first evaluation of the contribution of both factors: social isolation and epilepsy. We demonstrate that epilepsy in socially housed mice produces an activation of the HPA axis and anhedonia, responses which are amplified in isolated mice with epilepsy. A strong activation of the HPA axis in isolated animals with epilepsy may contribute to the expression of co-morbidities such as anxiety, cognitive deficits, and depression-like behavior.

The results obtained in mice are also worth discussing in a general manner. Animals were randomly selected at baseline for their future group assignment (treatment/housing). The first striking observation is how distributed are the values of anxiety and discrimination index of individual mice. The “control” population is far from homogeneous. As discussed above, this heterogeneity can be used to generate diverse groups of mice after an insult, e.g. mice susceptible and resilient to depression thus mimicking human conditions (Akil et al., 2018). Such heterogeneity is also found in rats, enabling to explain why some animals develop a severe form of epilepsy and/or co-morbidities and not others (Blugeot et al., 2011; Becker et al., 2015; Becker et al., 2019). Many factors can account for such heterogeneity in anxiety/cognitive performance at baseline in mice, including different genetic backgrounds and life experience (e.g. handling by the dam, dominant/dominated relations in the cage). Since we tested animals always at the same time of the day, circadian variations may not play a role. Multidien cycles (slower cycles with days-weeks-months periods) have been demonstrated in human (Baud et al., 2018) and rats (Baud et al., 2019) with epilepsy. Whether control animals display endogenous multidien cycles remains to be determined. If they do, anxiety/cognitive performance may vary as a function of these cycles, and since such slow cycles are not synchronized in animals recorded simultaneously (Baud et al., 2019), the anxiety/cognitive performance points will be distributed according to which position in the slow cycle a given animal is when it is tested.

Another striking observation we made is the linear relationship found between anxiety and ACTH/corticosterone/discrimination levels. The relationship was still present after isolation and Pilo treatment. This means that these variables are tied by a strong co-variance rule. It will be interesting to test other variables (e.g. stability of place cells, fear conditioning, working memory) with anxiety levels. Given the damage produced by pilocarpine-induced status epilepticus in numerous brain regions, we could have expected a deviation from the co-variance rule. Instead, animals just “slide down” the regression line towards stronger phenotypic expression. Said differently, epilepsy did not “reprogram” the relationship between anxiety and ACTH/corticosterone/discrimination levels. However, epilepsy did reprogram the anxiety/BDNF relationship. BDNF is involved in numerous functions and phenotypic traits, in particular, stress (Murinova et al., 2017) and epilepsy (Iughetti et al., 2018). Although there exists a linear relationship between anxiety and serum BNDF, the relationship is shifted towards larger anxiety value in Pilo mice, while the slope remains mostly unaffected. This suggests that epilepsy changes the general rule linking anxiety and serum BDNF levels. Together, our results further support the concept that rodents, like humans, should be considered as individuals. Even if as a group (on average) an effect can be evidenced, there are significant discrepancies between animals, including animals not responding as the group average. Again, many factors can account for such behavior, but we argue that discounting them may make us miss an important biological phenomenon that the observation revealed. It is therefore not surprising that following status epilepticus animals display different seizure severities.

In rats with epilepsy, we found that spontaneous seizures were, in average, 16 times more frequent in isolated animals as compared to animals kept in pairs or singly housed animals but with daily interactions with experimenter. Looking at individuals in the Isolated group, we noted a substantial dispersion of seizure frequency. Some Isolated animals had a low seizure frequency, in the range of the Handled/Paired groups (perhaps signing a resilience to stress-induced isolation or resilience to pilocarpine-induced status epilepticus). Even though, at baseline, animals may display a high variability (cf. the dispersion of ACTH, corticosterone and BDNF values in control rats), which may explain different fates as a function of housing conditions, the consequences of isolation are quite dramatic. During the three weeks of continuous recordings (starting six weeks after pilocarpine-induced status epilepticus), we did not find an increase in seizure frequency (Figure 9), suggesting that a global steady state activity had been reached (Williams et al., 2009). Interictal activity and seizures follow a multidien rhythm in the pilocarpine experimental rat model of epilepsy (Baud et al., 2019) as in humans (Baud et al., 2018). The fact that seizures occur at specific phases of this rhythm is not a confounding factor as the frequency of the multidien rhythm is between five and seven days in this rat model (Baud et al., 2019), which means that we captured at least three cycles.

In the present study, isolated animals had a high seizure frequency (≈16/day). It is difficult to compare this value with the existing literature as housing conditions are rarely mentioned. In addition, the beginning (how long after status epilepticus) and duration of 24/7 EEG recordings vary from one study to another. In the rat pilocarpine model, typical values for singly housed animals in Sprague Dawley rats are: 3/day during the early phase of epilepsy (Behr et al., 2017), 8/day (Paolone et al., 2019), 1-13/day (Tai et al., 2017), 10/day (Bankstahl et al., 2012); similar to values reported in Wistar rats: ≈4.5/day (Bajorat et al., 2011; Bajorat et al., 2018). Our study, although on the high end, remains in the same order of the magnitude as previously published work (we limited our analysis of the literature to the ongoing decade). We evaluated seizure severity only during the light phase, which constitutes a limitation of the study. We cannot rule out that seizures occurring during the dark phase may be less severe (including subclinical electrographic seizures). It is important to note that the International League Against Epilepsy released suggestions on common data elements for physiological (Gorter et al., 2018), behavioral (Mazarati et al., 2018) and EEG (Ono et al., 2018) studies performed in experimental animal models of epilepsy. Although our study was planned before the release of such information, adopting common data elements will help to compare different studies in the future.

Finding the mechanisms underlying the difference in phenotypes between the different housing conditions was beyond the scope of this study. We do not claim that the strong seizure phenotype found in the Isolated group is solely due to the stronger activation of the HPA axis, although it may contribute to it. We may tentatively propose that social isolation produces a vulnerability phenotype, as suggested by the low levels of serum BDNF found in Isolated animals (Becker et al., 2015). We suggest that epileptogenesis occurring in networks “weakened” by social isolation-induced stress would result in a strong phenotype.

The field of epilepsy research makes use of many different experimental models, strains and species. Results are model-, strain- and species-specific, which may explain some conflicting results. We propose that another confounding factor to consider is housing conditions. Some patients with epilepsy may experience social isolation, which correlates with more severe forms of epilepsy (Kotwas et al., 2017). Results obtained in experimental animal models kept in isolation may be highly relevant to such patient situation, or any other situation associated with high levels of stress. Results obtained in socially housed animals may be relevant to other subsets of patients, with a less severe phenotype. Our results have significant consequences in terms of data interpretation, in particular when testing anti-epileptogenesis or anti-seizure strategies. Our data provide strong evidence that the very large effect of Pilo under IC is an effect of Pilo *and* rearing. This means that the phenotype measured in singly housed animals is a blend of the treatment and the housing. It also means that any attempts to modify/treat epilepsy in these conditions could be modifying/treating the epilepsy effect or the housing effect, or both, because they are conflated. Some studies report a significant decrease in seizure frequency when animals are treated with a given compound. It is possible that the compound may have targeted the isolation-dependent stress component of the phenotype and not epilepsy itself. This is analogous to the anxiety/cognitive performance linear relationship we described. The two variables, housing and treatment, make the animals move up/down along the regression line. If the treatment affects the stress-induced component, animals will move back up to fewer seizures, but the biological component of epilepsy itself may not be affected *per se*. A reduction in seizure frequency by a factor of 16 would be equivalent to the difference we report between Paired and Isolated animals. For example, inhibition of kelch like ECH associated protein 1 (Keap1) reduces seizure frequency by a factor 20 in isolated kainic acid-treated Sprague-Dawley animals (Shekh-Ahmad et al., 2018). Keap1 controls nuclear factor erythroid 2-related factor 2 (Nrf2), a key regulator of antioxidant defense mechanisms. The authors suggest that these results support “the hypothesis that reactive oxygen species generation is a key event in the development of epilepsy” (Shekh-Ahmad et al., 2018). Since social isolation produces reactive oxygen species (Filipovic et al., 2017), it is possible that activating antioxidant defense mechanisms may have targeted the isolation-induced component of the seizure phenotype. Indeed, treating animals with a potent antioxidant did not affect seizure frequency in the kainic acid rat model when animals interacted daily with experimenters, while the treatment was very efficient when animals were characterized by high levels of reactive oxygen species before status epilepticus (Becker et al., 2019). These considerations provide another possible interpretation of the Keap1 results (Shekh-Ahmad et al., 2018). However, the conclusion of Shekh-Ahmad and collaborators that antioxidant treatment is strongly disease modifying in the context of social isolation is not only valid but also highly clinically relevant to socially isolated patients. Finally, many (often unpublished) preclinical studies performed in isolated animals did not evidence any significant effect on seizure frequency. Since the social isolation stress component could act as a confounding factor producing a strong phenotype, there may be many false negative results; i.e. the tested drugs may have been very efficient in experimental animals maintaining social interaction.

Although we cannot claim that maintaining social interaction between rodents in laboratory conditions corresponds to “normality” (as in the wild), single housing does produce strong biological alterations that render data interpretation more complex, in both physiological and pathological conditions. We recommend that material and methods should systematically report housing conditions and that all efforts should be made to maintain social interaction. Finally, these results were obtained in one experimental model (pilocarpine), in two species, one strain for each species and only in males. Similar studies should be performed in females, other strains and models such as kainic acid, kindling, tetanus toxin etc. Our results and conclusions should not be seen as a criticism of previous works. Rather varying housing conditions provide a unique opportunity to study, with the same experimental model, different situations found in patients with high and low stress levels.

## Acknowledgments

This work was supported by the European Union’s Seventh Framework Programme (FP7/2007 −2013) under grant agreement no602102 (EPITARGET), and Fondation pour la Recherche sur le Cerveau. HM was supported by NEUREN Project (PIRSES-GA −2012-318997). We thank Prof. Robert Calin-Jageman who helped us transitioning to the estimation based on confidence intervals.

## REFERENCES

1. Akil H, Gordon J, Hen R, Javitch J, Mayberg H, McEwen B, Meaney MJ, Nestler EJ (2018) Treatment resistant depression: A multi-scale, systems biology approach. Neurosci Biobehav Rev 84:272–288.

2. Amiri S, Haj-Mirzaian A, Amini-Khoei H, Razmi A, Shirzadian A, Rahimi-Balaei M, Olson CO, Mohsenzadeh A, Rastegar M, Zarrindast MR, Ghazi-Khansari M (2017) Protective effects of gabapentin against the seizure susceptibility and comorbid behavioral abnormalities in the early socially isolated mice. Eur J Pharmacol 797:106–114.

3. Arakawa H (2018) Ethological approach to social isolation effects in behavioral studies of laboratory rodents. Behav Brain Res 341:98–108.

4. Bajorat R, Wilde M, Sellmann T, Kirschstein T, Kohling R (2011) Seizure frequency in pilocarpine-treated rats is independent of circadian rhythm. Epilepsia 52:e118–122.

5. Bajorat R, Porath K, Kuhn J, Gossla E, Goerss D, Sellmann T, Kohling R, Kirschstein T (2018) Oral administration of the casein kinase 2 inhibitor TBB leads to persistent KCa2.2 channel up-regulation in the epileptic CA1 area and cortex, but lacks anti-seizure efficacy in the pilocarpine epilepsy model. Epilepsy Res 147:42–50.

6. Baldin E, Hauser WA, Pack A, Hesdorffer DC (2017) Stress is associated with an increased risk of recurrent seizures in adults. Epilepsia 58:1037–1046.

7. Bankstahl M, Bankstahl JP, Loscher W (2012) Inter-individual variation in the anticonvulsant effect of phenobarbital in the pilocarpine rat model of temporal lobe epilepsy. Exp Neurol 234:70–84.

8. Baud MO, Ghestem A, Benoliel JJ, Becker C, Bernard C (2019) Endogenous multidien rhythm of epilepsy in rats. Exp Neurol.

9. Baud MO, Kleen JK, Mirro EA, Andrechak JC, King-Stephens D, Chang EF, Rao VR (2018) Multi-day rhythms modulate seizure risk in epilepsy. Nat Commun 9:88.

10. Becker AJ (2018) Review: Animal models of acquired epilepsy: insights into mechanisms of human epileptogenesis. Neuropathol Appl Neurobiol 44:112–129.

11. Becker C, Bouvier E, Ghestem A, Siyoucef S, Claverie D, Camus F, Bartolomei F, Benoliel JJ, Bernard C (2015) Predicting and treating stress-induced vulnerability to epilepsy and depression. Ann Neurol 78:128–136.

12. Becker C, Mancic A, Ghestem A, Poillerat V, Claverie D, Bartolomei F, Brouillard F, Benoliel JJ, Bernard C (2019) Antioxidant treatment after epileptogenesis onset prevents comorbidities in rats sensitized by a past stressful event. Epilepsia 60:648–655.

13. Behr C, Levesque M, Stroh T, Avoli M (2017) Time-dependent evolution of seizures in a model of mesial temporal lobe epilepsy. Neurobiol Dis 106:205–213.

14. Bernard C (2016) The Diathesis-Epilepsy Model: How Past Events Impact the Development of Epilepsy and Comorbidities. Cold Spring Harb Perspect Med 6.

15. Berry A, Bellisario V, Capoccia S, Tirassa P, Calza A, Alleva E, Cirulli F (2012) Social deprivation stress is a triggering factor for the emergence of anxiety- and depression-like behaviours and leads to reduced brain BDNF levels in C57BL/6J mice. Psychoneuroendocrinology 37:762–772.

16. Bianchi M, Fone KF, Azmi N, Heidbreder CA, Hagan JJ, Marsden CA (2006) Isolation rearing induces recognition memory deficits accompanied by cytoskeletal alterations in rat hippocampus. Eur J Neurosci 24:2894–2902.

17. Blugeot A, Rivat C, Bouvier E, Molet J, Mouchard A, Zeau B, Bernard C, Benoliel JJ, Becker C (2011) Vulnerability to depression: from brain neuroplasticity to identification of biomarkers. J Neurosci 31:12889–12899.

18. Bouvier E, Brouillard F, Molet J, Claverie D, Cabungcal JH, Cresto N, Doligez N, Rivat C, Do KQ, Bernard C, Benoliel JJ, Becker C (2017) Nrf2-dependent persistent oxidative stress results in stress-induced vulnerability to depression. Mol Psychiatry 22:1795.

19. Brenes JC, Rodriguez O, Fornaguera J (2008) Differential effect of environment enrichment and social isolation on depressive-like behavior, spontaneous activity and serotonin and norepinephrine concentration in prefrontal cortex and ventral striatum. Pharmacol Biochem Behav 89:85–93.

20. Butler TR, Karkhanis AN, Jones SR, Weiner JL (2016) Adolescent Social Isolation as a Model of Heightened Vulnerability to Comorbid Alcoholism and Anxiety Disorders. Alcohol Clin Exp Res 40:1202–1214.

21. Chauviere L, Doublet T, Ghestem A, Siyoucef SS, Wendling F, Huys R, Jirsa V, Bartolomei F, Bernard C (2012) Changes in interictal spike features precede the onset of temporal lobe epilepsy. Ann Neurol 71:805–814.

22. Claverie D, Becker C, Ghestem A, Coutan M, Camus F, Bernard C, Benoliel JJ, Canini F (2016) Low beta2 Main Peak Frequency in the Electroencephalogram Signs Vulnerability to Depression. Front Neurosci 10:495.

23. Dezsi G, Ozturk E, Salzberg MR, Morris M, O’Brien TJ, Jones NC (2016) Environmental enrichment imparts disease-modifying and transgenerational effects on genetically-determined epilepsy and anxiety. Neurobiol Dis 93:129–136.

24. Djordjevic J, Djordjevic A, Adzic M, Radojcic MB (2012) Effects of chronic social isolation on Wistar rat behavior and brain plasticity markers. Neuropsychobiology 66:112–119.

25. El Houssaini K, Ivanov AI, Bernard C, Jirsa VK (2015) Seizures, refractory status epilepticus, and depolarization block as endogenous brain activities. Phys Rev E Stat Nonlin Soft Matter Phys 91:010701.

26. Ennaceur A, Aggleton JP (1994) Spontaneous recognition of object configurations in rats: effects of fornix lesions. Exp Brain Res 100:85–92.

27. Filipovic D, Todorovic N, Bernardi RE, Gass P (2017) Oxidative and nitrosative stress pathways in the brain of socially isolated adult male rats demonstrating depressive- and anxiety-like symptoms. Brain Struct Funct 222:1–20.

28. Gaskin S, Tardif M, Cole E, Piterkin P, Kayello L, Mumby DG (2010) Object familiarization and novel-object preference in rats. Behav Processes 83:61–71.

29. Gold PW (2015) The organization of the stress system and its dysregulation in depressive illness. Mol Psychiatry 20:32–47.

30. Gorter JA, van Vliet EA, Dedeurwaerdere S, Buchanan GF, Friedman D, Borges K, Grabenstatter H, Lukasiuk K, Scharfman HE, Nehlig A (2018) A companion to the preclinical common data elements for physiologic data in rodent epilepsy models. A report of the TASK3 Physiology Working Group of the ILAE/AES Joint Translational Task Force. Epilepsia Open 3:69–89.

31. Gunn BG, Baram TZ (2017) Stress and Seizures: Space, Time and Hippocampal Circuits. Trends Neurosci 40:667–679.

32. Han MH, Nestler EJ (2017) Neural Substrates of Depression and Resilience. Neurotherapeutics 14:677–686.

33. Hatch AM, Wiberg GS, Zawidzka Z, Cann M, Airth JM, Grice HC (1965) Isolation syndrome in the rat. Toxicol Appl Pharmacol 7:737–745.

34. Ieraci A, Mallei A, Popoli M (2016) Social Isolation Stress Induces Anxious-Depressive-Like Behavior and Alterations of Neuroplasticity-Related Genes in Adult Male Mice. Neural Plast 2016:6212983.

35. Inostroza M, Cid E, Menendez de la Prida L, Sandi C (2012) Different emotional disturbances in two experimental models of temporal lobe epilepsy in rats. PLoS One 7:e38959.

36. Inostroza M, Cid E, Brotons-Mas J, Gal B, Aivar P, Uzcategui YG, Sandi C, Menendez de la Prida L (2011) Hippocampal-dependent spatial memory in the water maze is preserved in an experimental model of temporal lobe epilepsy in rats. PLoS One 6:e22372.

37. Iughetti L, Lucaccioni L, Fugetto F, Predieri B, Berardi A, Ferrari F (2018) Brain-derived neurotrophic factor and epilepsy: a systematic review. Neuropeptides 72:23–29.

38. Jirsa VK, Stacey WC, Quilichini PP, Ivanov AI, Bernard C (2014) On the nature of seizure dynamics. Brain 137:2210–2230.

39. Kotloski RJ, Sutula TP (2015) Environmental enrichment: evidence for an unexpected therapeutic influence. Exp Neurol 264:121–126.

40. Kotwas I, McGonigal A, Bastien-Toniazzo M, Bartolomei F, Micoulaud-Franchi JA (2017) Stress regulation in drug-resistant epilepsy. Epilepsy Behav 71:39–50.

41. Krugel U, Fischer J, Bauer K, Sack U, Himmerich H (2014) The impact of social isolation on immunological parameters in rats. Arch Toxicol 88:853–855.

42. Lai MC, Holmes GL, Lee KH, Yang SN, Wang CA, Wu CL, Tiao MM, Hsieh CS, Lee CH, Huang LT (2006) Effect of neonatal isolation on outcome following neonatal seizures in rats--the role of corticosterone. Epilepsy Res 68:123–136.

43. Lapiz-Bluhm MD, Bondi CO, Doyen J, Rodriguez GA, Bedard-Arana T, Morilak DA (2008) Behavioural assays to model cognitive and affective dimensions of depression and anxiety in rats. J Neuroendocrinol 20:1115–1137.

44. Levesque M, Avoli M, Bernard C (2016) Animal models of temporal lobe epilepsy following systemic chemoconvulsant administration. Journal of neuroscience methods 260:45–52.

45. Lidster K, Jefferys JG, Blumcke I, Crunelli V, Flecknell P, Frenguelli BG, Gray WP, Kaminski R, Pitkanen A, Ragan I, Shah M, Simonato M, Trevelyan A, Volk H, Walker M, Yates N, Prescott MJ (2016) Opportunities for improving animal welfare in rodent models of epilepsy and seizures. J Neurosci Methods 260:2–25.

46. Loscher W (2017) Animal Models of Seizures and Epilepsy: Past, Present, and Future Role for the Discovery of Antiseizure Drugs. Neurochem Res 42:1873–1888.

47. Maguire J, Salpekar JA (2013) Stress, seizures, and hypothalamic-pituitary-adrenal axis targets for the treatment of epilepsy. Epilepsy Behav 26:352–362.

48. Mazarati A, Jones NC, Galanopoulou AS, Harte-Hargrove LC, Kalynchuk LE, Lenck-Santini PP, Medel-Matus JS, Nehlig A, de la Prida LM, Sarkisova K, Veliskova J (2018) A companion to the preclinical common data elements on neurobehavioral comorbidities of epilepsy: a report of the TASK3 behavior working group of the ILAE/AES Joint Translational Task Force. Epilepsia Open 3:24–52.

49. Mazarati AM, Shin D, Kwon YS, Bragin A, Pineda E, Tio D, Taylor AN, Sankar R (2009) Elevated plasma corticosterone level and depressive behavior in experimental temporal lobe epilepsy. Neurobiol Dis 34:457–461.

50. Medel-Matus JS, Shin D, Sankar R, Mazarati A (2017) Inherent vulnerabilities in monoaminergic pathways predict the emergence of depressive impairments in an animal model of chronic epilepsy. Epilepsia 58:e116–e121.

51. Moller M, Du Preez JL, Viljoen FP, Berk M, Emsley R, Harvey BH (2013) Social isolation rearing induces mitochondrial, immunological, neurochemical and behavioural deficits in rats, and is reversed by clozapine or N-acetyl cysteine. Brain Behav Immun 30:156–167.

52. Moran TP (2016) Anxiety and working memory capacity: A meta-analysis and narrative review. Psychol Bull 142:831–864.

53. Morelli E, Ghiglieri V, Pendolino V, Bagetta V, Pignataro A, Fejtova A, Costa C, Ammassari-Teule M, Gundelfinger ED, Picconi B, Calabresi P (2014) Environmental enrichment restores CA1 hippocampal LTP and reduces severity of seizures in epileptic mice. Exp Neurol 261:320–327.

54. Mumtaz F, Khan MI, Zubair M, Dehpour AR (2018) Neurobiology and consequences of social isolation stress in animal model-A comprehensive review. Biomed Pharmacother 105:1205–1222.

55. Murinova J, Hlavacova N, Chmelova M, Riecansky I (2017) The Evidence for Altered BDNF Expression in the Brain of Rats Reared or Housed in Social Isolation: A Systematic Review. Front Behav Neurosci 11:101.

56. Ngoupaye GT, Bum EN, Daniels WM (2013) Antidepressant-like effects of the aqueous macerate of the bulb of Gladiolus dalenii Van Geel (Iridaceae) in a rat model of epilepsy-associated depression. BMC Complement Altern Med 13:272.

57. Ono T, Wagenaar J, Giorgi FS, Fabera P, Hanaya R, Jefferys J, Moyer JT, Harte-Hargrove LC, Galanopoulou AS (2018) A companion to the preclinical common data elements and case report forms for rodent EEG studies. A report of the TASK3 EEG Working Group of the ILAE/AES Joint Translational Task Force. Epilepsia Open 3:90–103.

58. Paolone G, Falcicchia C, Lovisari F, Kokaia M, Bell WJ, Fradet T, Barbieri M, Wahlberg LU, Emerich DF, Simonato M (2019) Long-term, targeted delivery of GDNF from encapsulated cells is neuroprotective and reduces seizures in the pilocarpine model of epilepsy. J Neurosci.

59. Pellow S, Chopin P, File SE, Briley M (1985) Validation of open:closed arm entries in an elevated plus-maze as a measure of anxiety in the rat. J Neurosci Methods 14:149–167.

60. Rao RM, Sadananda M (2016) Influence of State and/or Trait Anxieties of Wistar Rats in an Anxiety Paradigm. Ann Neurosci 23:44–50.

61. Reger ML, Hovda DA, Giza CC (2009) Ontogeny of Rat Recognition Memory measured by the novel object recognition task. Dev Psychobiol 51:672–678.

62. Ros-Simo C, Valverde O (2012) Early-life social experiences in mice affect emotional behaviour and hypothalamic-pituitary-adrenal axis function. Pharmacol Biochem Behav 102:434–441.

63. Saggio ML, Spiegler A, Bernard C, Jirsa VK (2017) Fast-Slow Bursters in the Unfolding of a High Codimension Singularity and the Ultra-slow Transitions of Classes. J Math Neurosci 7:7.

64. Shao Y, Yan G, Xuan Y, Peng H, Huang QJ, Wu R, Xu H (2015) Chronic social isolation decreases glutamate and glutamine levels and induces oxidative stress in the rat hippocampus. Behav Brain Res 282:201–208.

65. Shekh-Ahmad T, Eckel R, Dayalan Naidu S, Higgins M, Yamamoto M, Dinkova-Kostova AT, Kovac S, Abramov AY, Walker MC (2018) KEAP1 inhibition is neuroprotective and suppresses the development of epilepsy. Brain 141:1390–1403.

66. Tai TY, Warner LN, Jones TD, Jung S, Concepcion FA, Skyrud DW, Fender J, Liu Y, Williams AD, Neumaier JF, D’Ambrosio R, Poolos NP (2017) Antiepileptic action of c-Jun N-terminal kinase (JNK) inhibition in an animal model of temporal lobe epilepsy. Neuroscience 349:35–47.

67. Veenema AH, Sijtsma B, Koolhaas JM, de Kloet ER (2005) The stress response to sensory contact in mice: genotype effect of the stimulus animal. Psychoneuroendocrinology 30:550–557.

68. Vrinda M, Sasidharan A, Aparna S, Srikumar BN, Kutty BM, Shankaranarayana Rao BS (2017) Enriched environment attenuates behavioral seizures and depression in chronic temporal lobe epilepsy. Epilepsia 58:1148–1158.

69. Williams PA, White AM, Clark S, Ferraro DJ, Swiercz W, Staley KJ, Dudek FE (2009) Development of spontaneous recurrent seizures after kainate-induced status epilepticus. JNeurosci 29:2103–2112.

70. Wulsin AC, Solomon MB, Privitera MD, Danzer SC, Herman JP (2016) Hypothalamic-pituitary-adrenocortical axis dysfunction in epilepsy. Physiol Behav 166:22–31.

